# Crosstalk between thrombospondin-1 and CD36 modulates platelet–RBC interaction limiting thrombosis and abdominal aneurysm formation

**DOI:** 10.1101/2024.03.18.585356

**Authors:** Kim Jürgen Krott, Tobias Feige, Agnes Bosbach, Irena Krüger, Susanne Pfeiler, Alicia Noeme Beele, Friedrich Reusswig, Elena Schickentanz-Dey, Alexandra Chadt, Malte Kelm, Norbert Gerdes, Kerstin Jurk, Klytaimnistra Kiouptsi, Christoph Reinhardt, Hadi Al-Hasani, Beate E. Kehrel, Saoussen Karray, Hubert Schelzig, Markus Udo Wagenhäuser, Margitta Elvers

**Author notes:** Correspondence to: Margitta Elvers, Ph.D., Clinic of Vascular and Endovascular Surgery, University Hospital Düsseldorf, Moorenstraße 5, 40225 Düsseldorf, Germany. Authorship note: MUW and ME are shared last authorship.

## Abstract

Red blood cells (RBCs) contribute to hemostasis and thrombosis by interaction with platelets via the FasL–FasR pathway to induce procoagulant activity and thrombin formation. Here, we identified a novel mechanism of platelet-RBC interaction via the CD36–thrombospondin-1 (TSP-1) signaling pathway, which is important in thrombus formation and the recruitment of RBCs to collagen-adherent platelets. Platelet-released TSP-1 can bind to CD36 at the RBC membrane to enhance procoagulant activity and to increase the activation of integrin α_IIb_β_3_, which represents an additional ligand for erythroid FasR, suggesting that both mechanisms of platelet–RBC interaction act in concert to propagate thrombus formation. In patients with abdominal aortic aneurysm (AAA), enhanced procoagulant activity of RBCs and platelets is accompanied by elevated exposure of TSP-1 and FasL at the platelet surface and accumulation of TSP-1 in the aortic wall and the intraluminal thrombus, suggesting that platelet–RBC interaction plays an important role in AAA pathology. TSP-1-deficient mice are protected against aortic diameter expansion in an experimental model of AAA, highlighting the crucial role of the CD36–TSP-1 axis in AAA. Thus, interfering with platelet–RBC interaction may be a promising therapeutic approach to reduce pro-coagulant activity and preserve AAA patients from surgery or rupture.

## INTRODUCTION

Platelets are important mediators of hemostasis. They not only prevent excessive blood loss at sites of vascular injury (1, 2) but also are central effectors in arterial thrombosis in myocardial infarction or stroke, which is the leading cause of death in Western countries (3, 4). Injury of the endothelium leads to the exposure of adhesive macromolecules from the extracellular matrix (ECM), for example collagen, fibronectin, laminin, von Willebrand factor (vWF), and TSP-1. These macromolecules serve as substrates to initiate the adhesion and activation of circulating platelets by the engagement of a wide spectrum of receptors such as glycoprotein (GP) Ib, GPVI, integrins, or CD36 (2). Platelet recruitment to the vessel wall and initial adhesion is dependent on the interaction between the platelet specific GPIb-V-IX complex and vWF (5). Collagens bind to the glycoprotein receptor GPVI and integrin α_2_β_1_, fibronectin engages both integrin α_5_β_1_ and laminin α_6_β_1_, and TSP-1 bind to CD36, among other receptors (6, 7). The secretion of TSP-1 from platelet α-granules and the interaction with CD36 induces an additional autocrine feed-forward loop that enforces the interaction of platelets in a growing thrombus and is important for thrombus stability (8, 9). During thrombus formation, TSP-1 protects vWF from cleavage through a disintegrin and metalloproteinase with a thrombospondin type 1 motif, member 13 (ADAMTS13), thus enabling the recruitment of circulating platelets to the growing thrombus by binding to GPIb (10, 11). Primary hemostasis is essential for initial coverage of injured vessels but is not sufficient for thrombus growth and stability (6). Thus, platelet activation is followed by blood coagulation to support the formation of stable arterial thrombi. Platelets are key factors in these processes because they provide a procoagulant surface by exposing the anionic phospholipid phosphatidylserine (PS) to allow the assembly of coagulation complexes on their membrane; this process is important for thrombin generation, fibrin formation and clot retraction (12).

In addition to the essential role of platelets, red blood cells (RBCs) are also known to support thrombus formation and stabilization. RBCs influence rheology in a passive manner (13) but also contribute to thrombus formation through the supply of ADP and ATP (14–17). Moreover, RBCs are able to expose PS on their membrane and thus contribute to the generation of thrombin (18–20). Recently, the mechanism by which RBCs influence thrombus formation by the exposure of PS has been identified (21). A small population of RBCs was identified in platelet thrombi and found to interact directly with platelets via the Fas ligand (FasL)– Fas receptor (FasR/CD95) pathway. In the presence of RBCs, FasL is exposed on the platelet surface to induce the exposure of PS at the RBC membrane by activation of the erythroid FasR, thereby enhancing thrombin generation. Inhibition or loss of either FasL or FasR results in decreased PS exposure of RBCs and reduced thrombin generation, leading to protection against venous and arterial thrombosis in mice (21). Other than FasL–FasR-mediated interactions, the mechanisms and consequences of platelet–RBC interaction for arterial thrombosis have not been well defined to date. Thus, the present study aimed to explore relevant mechanisms of platelet–RBC interactions. Here, we identified a new mechanism of platelets to activate RBCs via the release of TSP-1 that binds to CD36 at the RBC membrane to support PS exposure, which is critical for platelet activation in arterial thrombosis and the pathology of abdominal aortic aneurysm (AAA).

## RESULTS

### Erythroid CD36 supports PS exposure of red blood cells (RBCs) and thrombus formation on collagen under arterial shear rates *ex vivo*

CD36 is exposed on the membrane of human platelets (Figure 1A) and has been shown to play an important role in thrombus formation (8, 9, 22, 23). Similarly, RBCs expose CD36 at their membrane (Figure 1A), but its role on erythroid cells has not been described. To investigate whether CD36 on RBCs is involved in platelet–RBC interactions and procoagulant activity, we determined annexin V binding of human platelets (Figure 1B) and RBCs (Figure 1C) that had been treated with a blocking anti-CD36 antibody under static conditions using flow cytometry. Blocking of CD36 only at the RBC membrane resulted in reduced annexin V binding of platelets while treatment of platelets did not alter PS exposure (Figure 1B). In contrast, concomitant blocking of CD36 on both, collagen-related peptide (CRP)-activated platelets and RBCs resulted in reduced PS exposure on RBCs, suggesting that CD36 plays a role in platelet–RBC interactions (Figure 1C).

**Figure 1.**
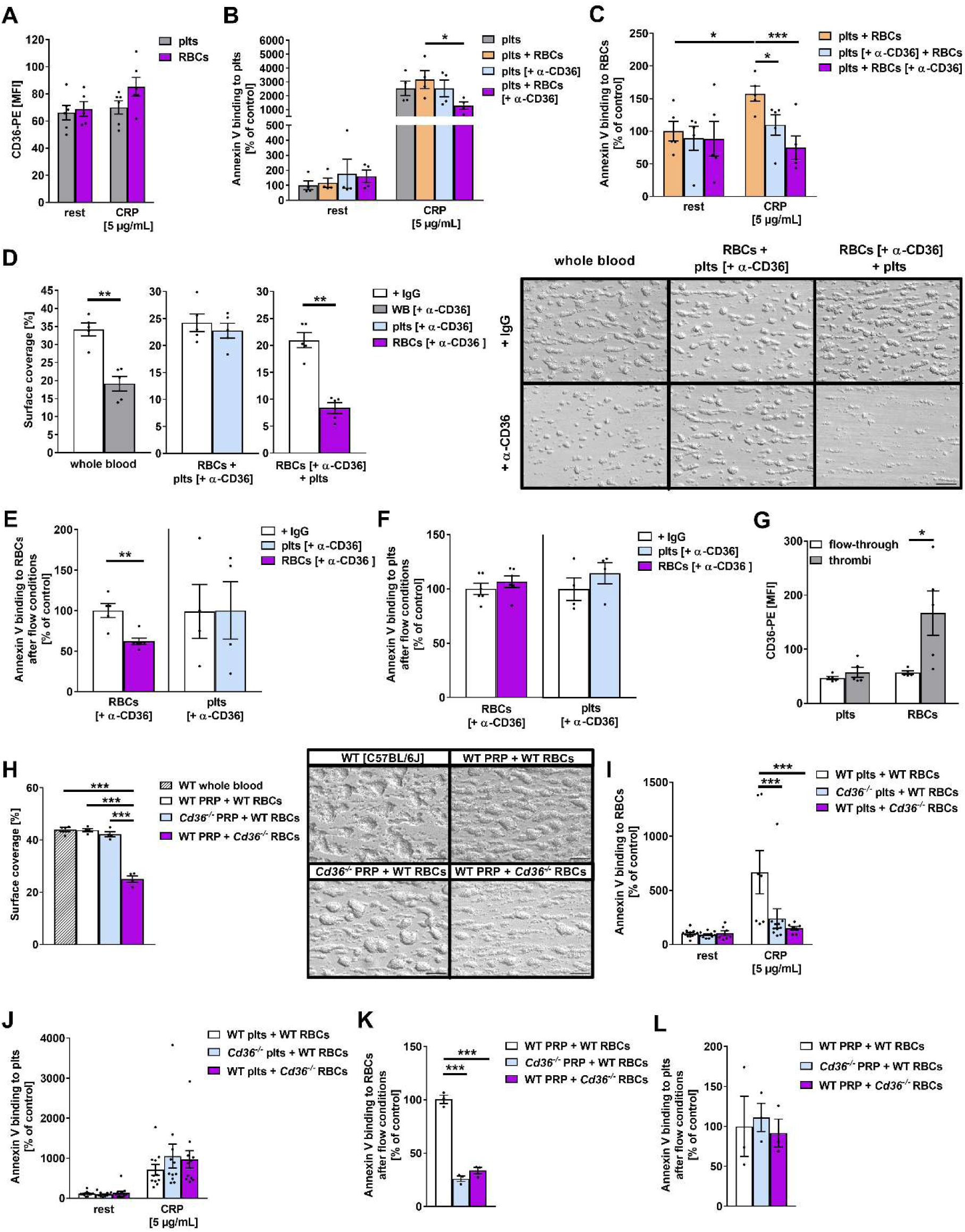
Erythroid CD36 supports thrombus formation on collagen under arterial shear rates. (**A**) CD36 externalization on human platelets and RBCs (*n* = 6). (**B** and **C**) annexin V binding to resting and CRP-stimulated human platelets and RBCs under static conditions determined by flow cytometry (*n* = 4–5), indicated as percent-gated cells normalized to controls. Cells were pre-incubated independently with an anti-CD36 mAb (2 μg/mL) and RBCs were subsequently washed before co-incubation with platelets. (**D**) Surface coverage of thrombus formation on collagen (200 µg/mL) at a shear rate of 1,700 s^−1^. Human platelets, RBCs and plasma were isolated and either RBCs or platelets as well as whole blood were incubated with an anti-CD36 mAb (2 µg/mL, RBCs, purple; plts, light blue). RBCs were washed before reassembling with platelets and plasma (*n* = 5). IgG served as control (white pillar). Representative differential interference contrast (DIC) images are shown. Scale bar = 50 µm. (**E** and **F**) Annexin V binding to human RBCs and platelets isolated from thrombi after flow chamber experiments at a shear rate of 1,700 s^−1^ with or without an anti-CD36 mAb treatment (2 µg/mL, RBCs, purple; plts, light blue) (*n* = 4–6). IgG served as control (white pillar). (**G**) CD36 externalization of human platelets and RBCs isolated from thrombi and flow-through (*n* = 5). (**H**) Surface coverage of thrombus formation on collagen (200 µg/mL) at a shear rate of 1,700 s^−1^. Either WT (C57BL/6J) or *Cd36*^−/–^ PRP was supplemented with either WT RBCs or *Cd36*^−/–^ RBCs (*n* = 4). Whole blood samples of C57BL/6J mice served as controls (*n* = 4). Representative DIC images are shown. Scale bar = 50 µm. (**I**–**L**) Annexin V binding to RBCs or platelets under static and dynamic conditions determined by flow cytometry. Either WT or *Cd36*^−/–^ RBCs were reassembled with either WT or *Cd36*^−/–^ platelets (**I** and **J**) or PRP (**K** and **L**) (*n* = 3–12). Data are represented as mean values ± SEM. \**P* < 0.05; \*\**P* < 0.01; \*\*\**P* < 0.001 tested by two-way ANOVA (**A**–**C**), student’s t-test (**D**–**F** and **H**) and one-way ANOVA with Sidak’s multiple comparison test (**G**–**L**). CRP, collagen-related peptide; rest, resting; plts, platelets; RBC, red blood cells; PRP, platelet-rich plasma; WT, wild-type; MFI, mean fluorescence intensity.

Next, we analyzed platelet adhesion and thrombus formation under flow using a flow chamber system to examine whether CD36 on RBCs plays a role in thrombus formation. To this end, we used either whole blood or platelet-rich plasma (PRP) that had been supplemented with RBCs (4 × 10^6^ RBCs/µL, corresponding to a physiological hematocrit of 40%). CD36 was blocked by antibody treatment of whole blood (Figure 1D, left panel) only on the surface of platelets (Figure 1D, middle panel) or only on the RBC membrane (Figure 1D, right panel). As shown in Figure 1D, antibody treatment of whole blood or restricted to RBCs led to reduced thrombus formation, whereas no differences in thrombus formation were observed when only platelet CD36 was blocked (Figure 1D). Reduced thrombus formation from blocking CD36 at the RBC membrane was accompanied by reduced PS exposure of RBCs (Figure 1E) but unaltered PS exposure of platelets (Figure 1F) after we isolated these cells from the thrombi by Accutase^®^ treatment and determined annexin V binding by flow cytometry. Blocking of platelet CD36 did not alter annexin V binding of either RBCs (Figure 1E) or platelets (Figure 1F). In contrast to static experiments showing unaltered CD36 exposure of platelets and RBCs under resting and CRP-stimulated conditions (Figure 1A), we detected significantly enhanced levels of CD36 on the membrane of RBCs that had been isolated from the thrombi as compared to RBCs isolated from the flow-through (Figure 1G). However, CD36 expression at the platelet membrane was not altered between platelets isolated from the flow-through and those from thrombi.

To confirm the functional relevance of erythroid CD36 for PS exposure and thrombus formation, we next used CD36-deficient (*Cd36*^−/–^) mice. Experiments using whole blood revealed reduced collagen-dependent thrombus formation under a shear rate of 1,700 s^−1^ but no differences at a shear rate of 1,000 s^−1^, in line with previous finings (Supplemental Figure 1A and B) (8). Next, we isolated PRP and RBCs from CD36-deficient mice. We detected reduced thrombus formation under flow conditions only when PRP from wild-type (WT) mice and RBCs from CD36-deficient mice were perfused through the chamber (Figure 1H). In contrast, no alterations in thrombus formation were observed with whole blood from control mice (C57BL/6J), PRP, and RBCs from WT mice or when PRP from CD36-deficient mice and WT RBCs were perfused through the chamber. These results clearly indicate that CD36 on RBCs plays an important role in thrombus formation under flow (Figure 1H). In line with reduced thrombus formation, we detected reduced FasL exposure of platelets in the presence of CD36-deficient RBCs while FasL exposure of platelets in PRP from CD36-deficient mice in the presence of WT RBCs was unaltered compared to controls (platelets and RBCs from WT mice, Supplemental Figure 1C). These results suggest that erythroid CD36 is not only involved in the interaction of platelets and RBCs but also promotes FasL–FasR-mediated signaling that represents an important mechanism of platelet-induced RBC activation to facilitate thrombus formation by enhanced procoagulant activity (21). Furthermore, CD36 deficiency of both platelets and RBCs led to significantly reduced PS exposure of RBCs compared to platelets and RBCs from WT mice under static conditions (Figure 1I), confirming the results with human blood cells (Figure 1C). However, annexin V binding to platelets was not altered, for either CD36-deficient platelets or CD36-deficient RBCs (Figure 1J). PS exposure of murine platelets and RBCs was also analyzed under flow conditions (Figure 1K and L). Again, annexin V binding to RBCs was significantly reduced but no alterations in PS exposure of platelets were detected when either CD36-deficient platelets or RBCs were used (Figure 1K and L).

### Thrombospondin-1 binds to erythroid CD36 to support platelet adhesion and thrombus formation

To characterize the impact of erythroid CD36 in thrombus formation and the underlying mechanisms, we performed flow chamber experiments using different matrices, including TSP-1, collagen, and collagen–TSP-1 at a shear rate of 1,700 s^−1^. Therefore, PRP and RBCs from healthy volunteers were perfused through the flow chamber. Although no thrombus formation was observed under flow conditions on a TSP-1 matrix, we detected elevated thrombus formation on an immobilized collagen–TSP-1 matrix compared to collagen alone (Figure 2A). Inhibition of CD36 only at the RBC membrane significantly reduced thrombus formation on collagen and on collagen–TSP-1 compared to controls. Furthermore, thrombus formation was found to be significantly reduced after comparing the surface coverage of three-dimensional thrombi formed on the collagen matrix compared to the collagen–TSP-1 matrix when CD36 was blocked on RBCs (Figure 2A). In contrast, the inhibition of CD36 on platelets did not alter thrombus formation on collagen, but on a collagen–TSP-1 matrix under flow conditions using a shear rate of 1,700 s^−1^ (Figure 2B). Using a blocking anti-TSP-1 antibody, we could confirm the importance of TSP-1 for thrombus formation under flow by detecting significantly reduced surface coverage of thrombi compared to IgG-treated controls (Figure 2C).

**Figure 2.**
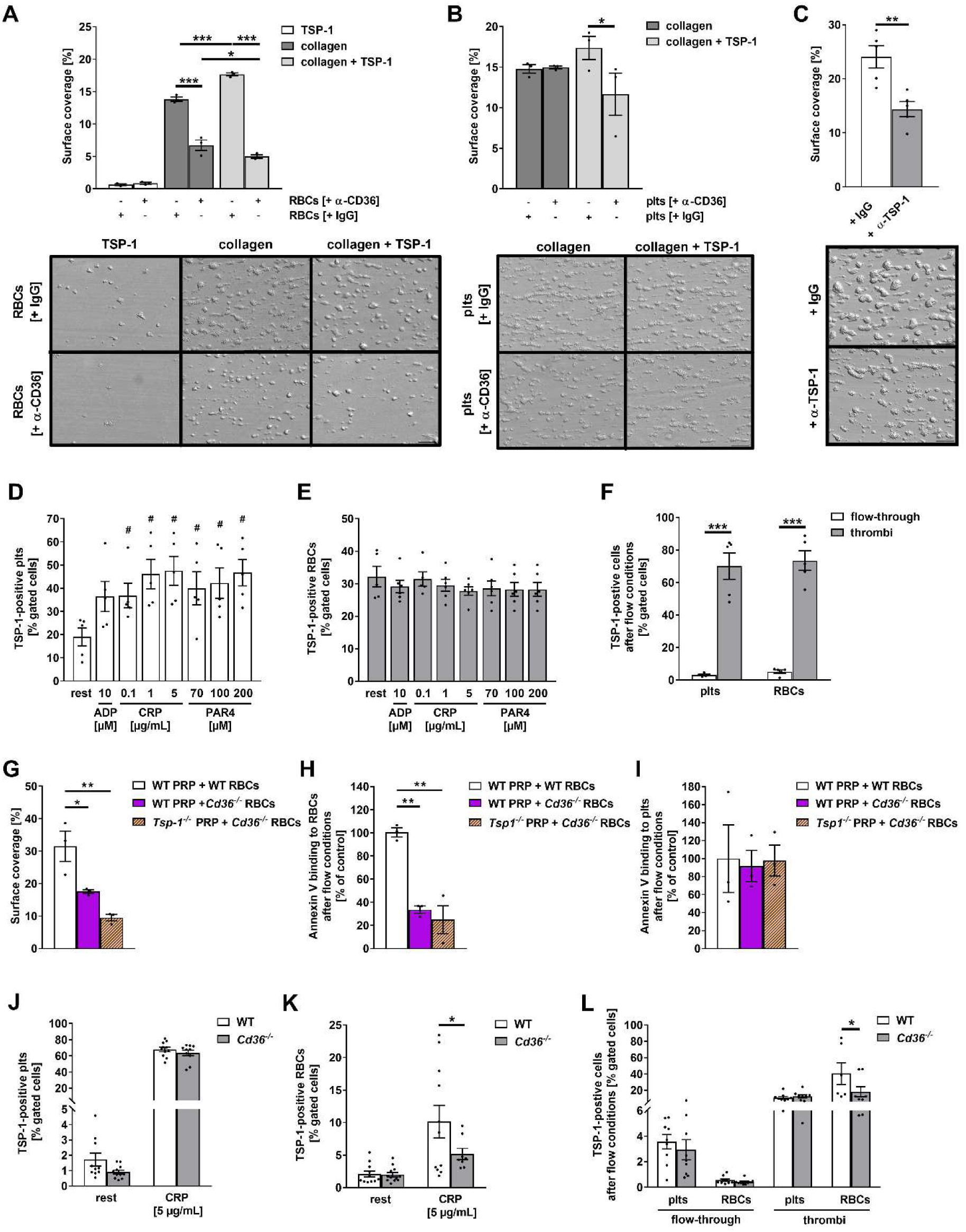
Platelet-released TSP-1 and CD36 are important for platelet adhesion, procoagulant surface of RBCs and thrombus formation. (**A** and **B**) Surface coverage of thrombus formation on a TSP-1 (100 µg/mL), collagen (200 µg/mL) or collagen–TSP-1 matrix at a shear rate of 1,700 s^−1^. Human platelets, RBCs and plasma were isolated and either RBCs (**A**) or platelets (**B**) were incubated with a without an anti-CD36 mAb (2 µg/mL). Subsequently, RBCs were washed before reassembling with platelets and plasma (*n* = 3). Representative DIC images are shown. Scale bar: 50 µm. (**C**) Surface coverage of thrombus formation on a collagen matrix at a shear rate of 1,700 s^−1^ of human whole blood preincubated with or without anit-TSP-1 mAb (2 µg/mL) (*n* = 5). Representative DIC images are shown. Scale bar: 50 µm. (**D** and **E**) Flow cytometric analysis of TSP-1 binding to platelets (**D**) or RBCs (**E**) in human whole blood upon stimulation with indicated agonists (*n* = 6). (**F**) Flow cytometric analysis of TSP-1 binding to human platelets and RBCs isolated from thrombi and flow-through after thrombus formation at a shear rate of 1,700 s^−1^ (*n* = 5). (**G**) Surface coverage of thrombus formation on a collagen matrix (200 µg/mL) at a shear rate of 1,700 s^−1^. Either WT or *Cd36*^−/–^ RBCs were reassembled with either WT or *Tsp-1*^−/–^ PRP (*n* = 3). (**H** and **I**) Annexin V binding to RBCs (**H**) or platelets (**I**) after flow chamber experiments shown in figure 2G (*n* = 3). (**J** and **K**) TSP-1 binding to platelets (**J**) and RBCs (**K**) in whole blood of WT and *Cd36*^−/–^ mice (*n* = 10). (**L**) TSP-1 binding to either WT or *Cd36*^−/–^ platelets and RBCs isolated from thrombi after flow chamber experiments at a shear rate of 1,700 s^−1^ (*n* = 9). Data are represented as mean values ± SEM. \**P* < 0.05; \*\**P* < 0.01; \*\*\**P* < 0.001; # indicates significant analysis vs. resting conditions tested by two-way ANOVA (**A, B, F,** and **J**–**L**), student’s t-test (**C**) and one-way ANOVA with Sidak’s multiple comparison test (**D, E** and **G**–**I**). Plts, platelets; RBC, red blood cells; WT, wild-type; WB, whole blood; rest, resting; ADP, adenosine diphosphate; CRP, collagen-related peptide; PAR4, protease-activated receptor 4 activating peptide.

Platelets possess a high content of TSP-1 protein, which can be released upon activation; TSP-1 is not present in RBCs (Supplemental Figure 2). Therefore, we measured TSP-1 binding to platelets (Figure 2D) and RBCs (Figure 2E) under static conditions using whole blood from healthy volunteers. Platelets were activated with indicated agonists and TSP-1-positive cells were determined by flow cytometry (Figure 2D and E). We detected binding of the anti-TSP-1 antibody to platelets and RBCs. While TSP-1 binding to platelets (Figure 2D) was increased upon platelet activation, we did not observe alterations in TSP-1 binding to RBCs in the presence of resting or activated platelets using different agonists (Figure 2E). The presence of platelets was sufficient to induce TSP-1 binding to RBCs. In contrast, TSP-1 binding to both platelets and RBCs was massively increased when we isolated these cells from collagen-adherent thrombi by Accutase^®^ treatment following flow chamber experiments (Figure 2F), suggesting that flow conditions and the incorporation of platelets and RBCs into the growing thrombus affect TSP-1 binding to these cells.

We next confirmed the results obtained with human platelets using platelets from WT, CD36, and TSP-1-deficient mice. First, analysis of thrombus formation under flow conditions revealed significantly reduced formation of three-dimensional thrombi on a collagen matrix using either WT or TSP-1-deficient PRP incubated with either WT or CD36-deficient RBCs (Figure 2G, Supplemental Figure 1D). PS exposure at the surface of RBCs (Figure 2H) and platelets (Figure 2I) isolated from these thrombi revealed no differences in the number of annexin V-positive platelets, while the number of procoagulant RBCs was reduced when CD36-deficient RBCs and WT or TSP-1-deficient platelets were used for experiments (Figure 2H and I). Thus, these data provide the first evidence for an interaction of TSP-1 with erythroid CD36 that is important in thrombus formation under arterial shear.

To confirm TSP-1 binding to erythroid CD36 upon murine platelet activation, we determined TSP-1 binding to RBCs and platelets under static (Figure 2J and K) and flow conditions (Figure 2L) using whole blood from CD36-deficient mice. In static experiments, platelets were activated with CRP, and TSP-1-positive RBCs and platelets were determined by flow cytometry (Figure 2J and K). Under these conditions, TSP-1 binding was unaltered between CD36-deficient and WT control platelets. In contrast, the binding of TSP-1 to CD36-deficient RBCs was significantly reduced when platelets were activated with CRP compared to WT controls (Figure 2K). Under flow conditions, TSP-1 binding to CD36-deficient RBCs isolated from collagen-adherent thrombi was significantly reduced whereas no alterations were detected using CD36-deficient platelets compared to platelets from WT mice (Figure 2L). However, only limited binding of TSP-1 to platelets and RBCs from the flow-through was observed, confirming the results with human platelets and RBCs and demonstrating strongly increased binding of TSP-1 to platelets and RBCs in the thrombus but not in the flow-through.

### Platelet–RBC interactions via CD36–TSP-1 and FasL–FasR modulate integrin β_3_ externalization and platelet aggregation

In a further set of experiments, we investigated the impact of platelet–RBC interactions on integrin α_IIb_β_3_ exposure and activation. Flow cytometric analysis revealed increased exposure of integrin α_IIb_β_3_ at the surface of human platelets under resting and activated conditions when platelets were incubated with RBCs compared to platelets alone (Figure 3A). RBC-mediated upregulation of integrin α_IIb_β_3_ was at least in part dependent on erythroid CD36 (Figure 3B) but not on TSP-1 (Figure 3C). To analyze whether the interaction of platelets and RBCs via the FasL–FasR axis plays any role for integrin α_IIb_β_3_ at the platelet surface, we blocked either FasR or FasL on human RBCs and platelets (Supplemental Figure 3A and C) or used murine FasR-deficient RBCs or FasL-deficient platelets (Supplemental Figure 3B and D). Inhibition or deficiency of FasR at the RBC membrane revealed that RBC-mediated upregulation of integrin α_IIb_β_3_ depends on erythroid FasR in both humans and mice (Supplemental Figure 3A and B). However, blocking of FasL by treatment with hDcR3 (recombinant human decoy receptor 3 protein) or the use of FasL-deficient platelets did not modulate integrin α_IIb_β_3_ exposure at the surface of human or murine platelets (Supplemental Figure 3C and D).

**Figure 3.**
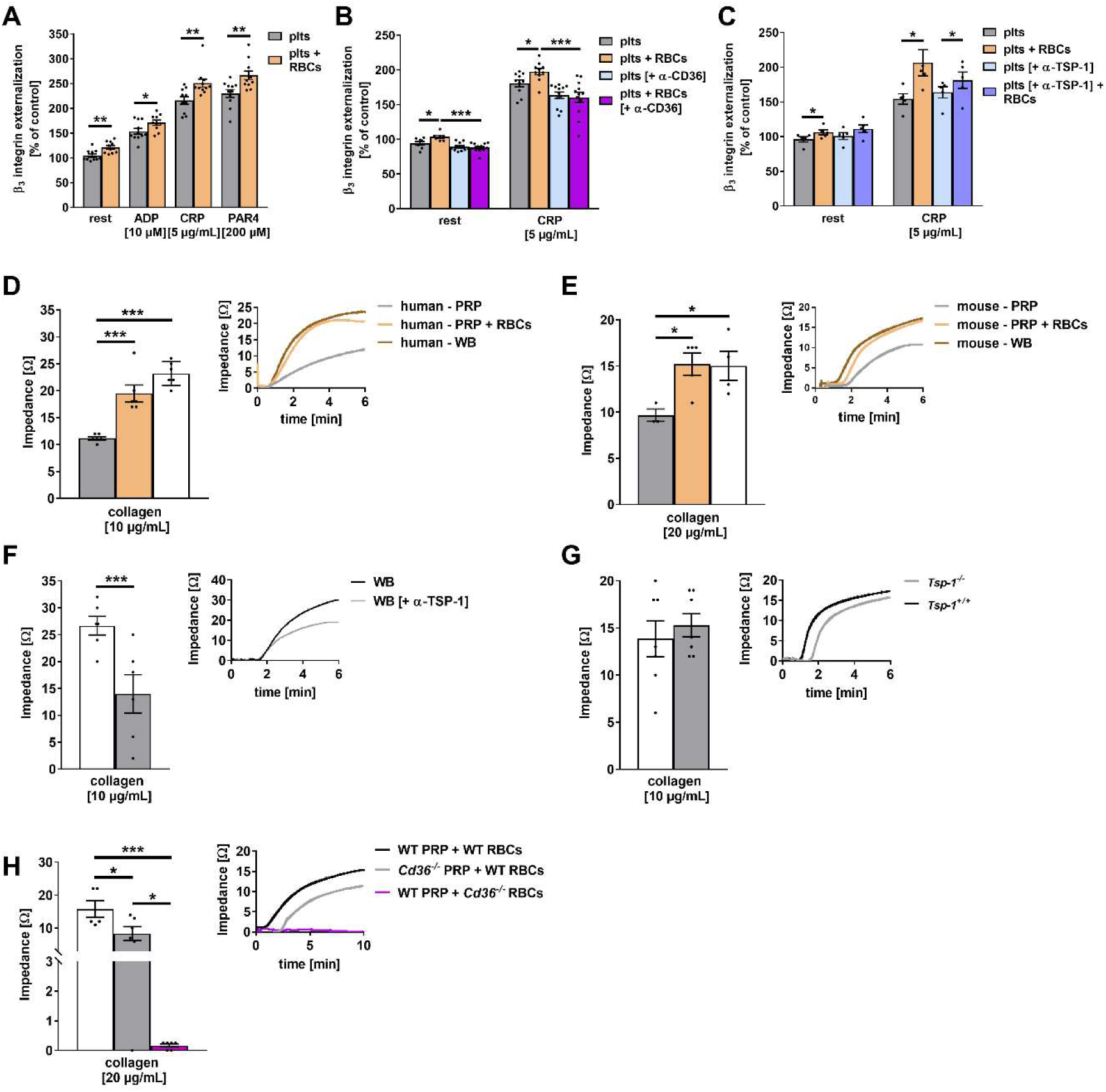
CD36–TSP-1 axis influences integrin β_3_ externalization and platelet aggregation *in vitro*. (**A–C**) Externalization of integrin β_3_ subunit (CD61-PE) on the surface of human platelets in the absence or presence of RBCs after stimulation with indicated agonists, determined by flow cytometry. (**A**) Integrin β_3_ of human platelets without antibody treatment (*n* = 11). (**B** and **C**) Either human isolated platelets (light blue) or RBCs (purple) were independently treated with an anti-CD36 mAb (**B**, 2 µg/mL, *n* = 10–12) or an anti-TSP-1 mAb (**C**, 2 µg/mL, *n* = 4–5). (**D**–**H**) Impedance measurements of human or murine whole blood and PRP in the absence or presence of RBCs after stimulation with collagen was performed. Representative aggregation curves are shown. (**D** and **E**) Isolated PRP and RBCs from human and mice were reassembled with platelet free plasma (*n* = 4–6). (**F**) Human whole blood with an anti-TSP-1 mAb treatment (2 µg/mL) after stimulation with collagen (10 µg/mL) (*n* = 6). (**G**) Whole blood samples of *Tsp-1*^−/–^ mice after stimulation with collagen (10 µg/mL) (*n* = 7). (**H**) Platelets from either WT or *Cd36*^−/–^ mice were incubated with either WT or *Cd36*^−/–^ RBCs after stimulation with collagen (20 µg/mL) (*n* = 5–6). Data are represented as mean values ± SEM. \**P* < 0.05; ** *P* < 0.01;*** *P* < 0.001 tested by two-way ANOVA (**A**–**C**), one-way ANOVA (**D, E** and **H**) with Sidak’s multiple comparison test and student’s t-test (**F** and **G**). Plts, platelets; PRP, platelet rich plasma; RBC, red blood cells; WT, wild-type; WB, whole blood; rest, resting; ADP, adenosine diphosphate; CRP, collagen-related peptide; PAR4, protease-activated receptor 4 activating peptide.

To analyze the contribution of integrin α_IIb_β_3_ to platelet–RBC interactions, we allowed human platelets to adhere to immobilized recombinant FasR under static conditions. Platelet activation with ADP significantly increased platelet adhesion to FasR (Supplemental Figure 3E and F). Treatment of platelets with the integrin α_IIb_β_3_ inhibitor tirofiban reduced platelet adhesion to recombinant FasR under ADP-stimulated but not under resting conditions (Supplemental Figure 3E). Reduced platelet adhesion to FasR was also observed when we blocked platelet FasL with hDcR3. In contrast to tirofiban treatment, blocking of FasL reduced platelet adhesion to FasR under resting as well as under ADP-stimulating conditions. However, we observed no additive effects of tirofiban and hDcR3 treatment on platelet adhesion to recombinant FasR. Also in contrast to tirofiban treatment, we detected reduced platelet adhesion to recombinant FasR under resting and ADP-stimulating conditions when we blocked integrin α_IIb_β_3_ using the antibody abciximab. Furthermore, an additive effect on platelet binding to recombinant FasR was observed when platelets were treated with abciximab and hDcR3 to block integrin α_IIb_β_3_ and FasL simultaneously at the platelets surface (Supplemental Figure 3F). This effect may have occurred because abciximab has strong affinity to its target and therefore might occupy the receptors for a long period, while tirofiban is a reversible antagonist of integrin α_IIb_β_3_. Binding of integrin α_IIb_β_3_ to immobilized FasR was further confirmed using CHO cells that had been stably transfected with human integrin α_IIb_β_3_ (A5 CHO, Supplemental Figure 3G). CHO and A5 CHO cells were allowed to adhere to immobilized FasR. A significant number of A5 CHO cells adhered to recombinant FasR, while CHO cells that did not express integrin α_IIb_β_3_ nearly failed to adhere to recombinant FasR. In control experiments, the adhesion of A5 CHO cells to immobilized fibrinogen (positive control) and BSA (negative control) was analyzed, showing strongly reduced ability of CHO cells to bind to these proteins compared to A5 CHO cells.

To investigate the impact of integrin α_IIb_β_3_ on PS exposure of RBCs and platelets, we used abciximab to block integrin α_IIb_β_3_ and hDcR3 to block FasL at the platelet surface (Supplemental Figure 3H and I). In the presence of the platelet agonists ADP or CRP, blocking of either integrin α_IIb_β_3_ or FasL led to reduced PS exposure on the surface of platelets (Supplemental Figure 3H) and RBCs (Supplemental Figure 3I); however, no additive effects on PS exposure of platelets and RBCs were observed. To confirm the impact of FasL and integrin α_IIb_β_3_ on PS exposure of platelets and RBCs, we performed experiments using mice with a platelet-specific deletion of FasL (*Fasl^fl/fl^-Pf4-Cre^+^*). In line with published data using blood from healthy volunteers (24), blocking of integrin α_IIb_β_3_ (with antibody Leo.H4) at the surface of murine platelets reduced PS exposure by these cells. Furthermore, inhibition of integrin α_IIb_β_3_ led to reduced PS exposure on the membrane of RBCs isolated from *Fasl^fl/fl^-Pf4-Cre*^−^ mice as well (Supplemental Figure 3J and K). Genetic deletion of FasL results in reduced PS exposure at the surface of platelets and RBCs (21). Blocking of integrin α_IIb_β_3_ at the surface of FasL-deficient platelets resulted in a further decrease of PS exposure at the surface of both platelets and RBCs (Supplemental Figure 3J and K). These results implicate two independent ligands by which platelet–RBC interactions induce procoagulant activity of RBCs via activation of erythroid FasR and erythroid CD36.

Platelet aggregation is mediated by integrin α_IIb_β_3_, which induces the formation of aggregates via binding to fibrinogen. To investigate whether RBCs exert an impact on platelet aggregation via integrin α_IIb_β_3_, we analyzed platelet aggregation using platelets alone (PRP), platelets supplemented with RBCs (PRP + RBCs), or whole blood from healthy volunteers or WT mice (Figure 3D and E). In response to collagen, platelet aggregation was significantly enhanced when whole blood or the combination of PRP + RBCs was used as compared to PRP alone. Inhibition of TSP-1 by antibody treatment of human whole blood resulted in reduced platelet aggregation (Figure 3F). In contrast, whole blood from TSP-1-deficient mice showed no differences compared to controls (Figure 3G), as shown previously by others (25). CD36 is involved in the regulation of platelet aggregation, as genetic deficiency of CD36 of either platelets or RBCs resulted in significantly reduced platelet aggregation following stimulation with collagen (Figure 3H). The interaction of platelets and RBCs via FasL–FasR modulates platelet aggregation as well, as the loss of either FasR or FasL led to reduced platelet aggregation in response to collagen using whole blood from FasR-deficient mice or mice with platelet-specific deficiency of FasL compared to control mice (Supplemental Figure 4A and B).

### Active recruitment of RBCs allows platelet–RBC interactions on collagen under flow

Various previous studies have provided strong evidence that RBCs support thrombus formation in hemostasis and thrombosis by direct and indirect mechanisms. However, the mechanisms by which RBCs are captured by platelets and which receptors are involved to incorporate RBCs into the growing thrombus remain elusive. Therefore, we analyzed the recruitment of RBCs to activated platelets to allow interaction of these cells upon thrombus formation. Platelets were allowed to adhere to a collagen matrix that had been immobilized on a coverslip (Figure 4A). This coverslip was then used in flow chamber experiments to investigate the capture of RBCs to these adhesive and activated platelets under flow conditions. To this end, RBCs were isolated and perfused through the chamber to allow contact with and recruitment by collagen-adherent platelets (Figure 4B). Under arterial shear rates (1,700 s^−1^), only transient adherent RBCs could be detected under these conditions. First, the impact of PS externalization on the membrane of platelets and RBCs was analyzed for its contribution to the active recruitment of RBCs. As shown in Figure 4C, annexin V treatment of both RBCs and platelets resulted in a strongly reduced number of transient adherent RBCs. To differentiate between cells, either platelets or RBCs were treated with annexin V and the number of transient adherent RBCs was determined. The results demonstrate that the exposure of PS on the platelet as well as on the RBC membrane is important for the recruitment of RBCs to collagen-adherent platelets (Figure 4D).

**Figure 4.**
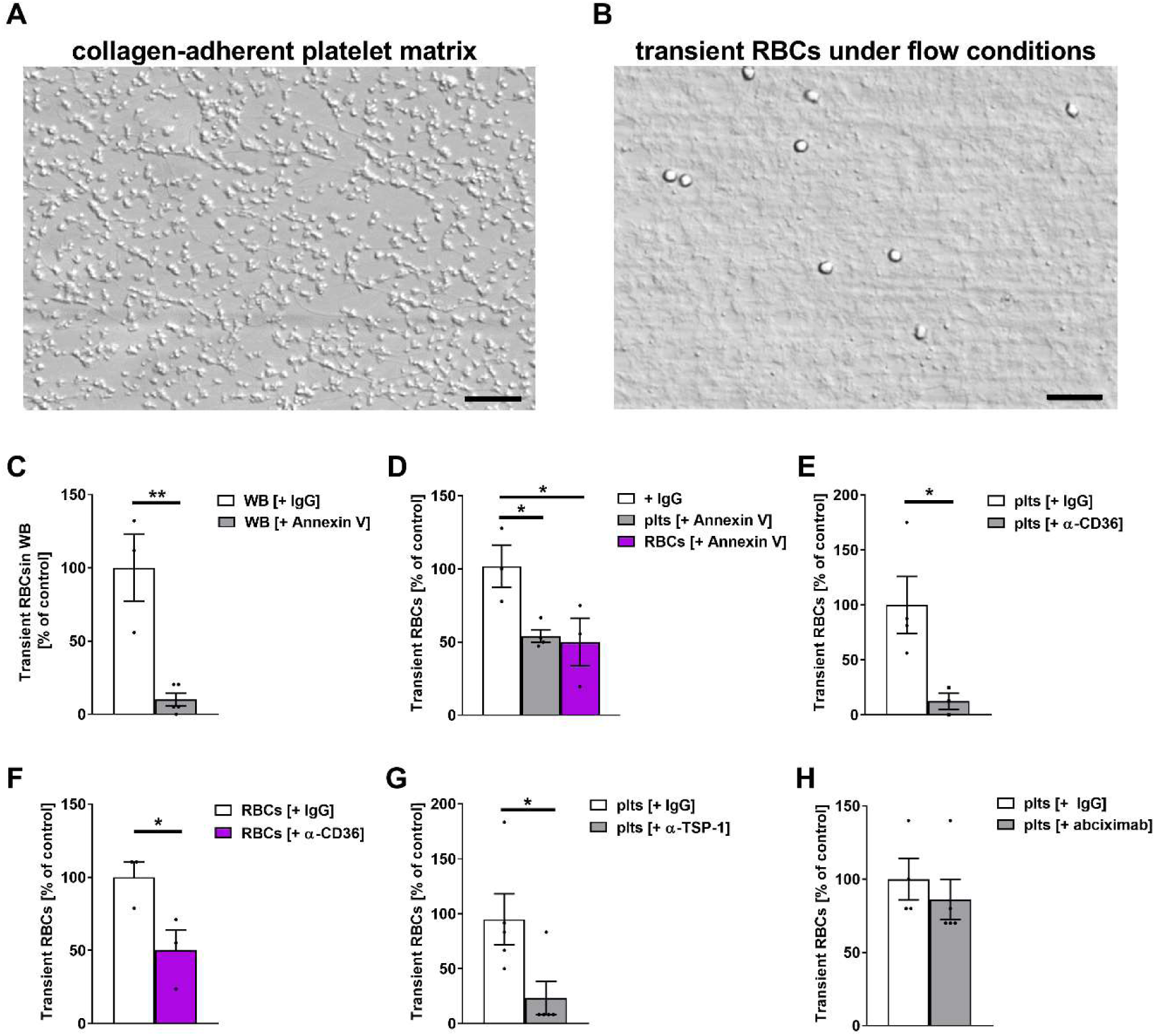
Active recruitment of RBCs to collagen-adherent platelets. (**A**) Representative image of the collagen-adherent platelet matrix. Isolated human platelets were stimulated with ADP (10 µM) and allowed to adhere on collagen (200 µg/mL) coated coverslips for 10 min. Scale bar: 50 µm. (**B**) Representative image of tethering RBCs on collagen-adherent platelets under flow using a shear rate of 1,700 s^−1^. Scale bar: 50 µm. (**C**) Both, platelets and RBCs, were incubated with recombinant annexin V inhibiting protein (5 µg/mL) before perfusion through the chamber and compared to untreated controls (*n* = 4). (**D**) Either platelets or RBCs were treated with recombinant annexin V inhibiting protein to block PS exposure (*n* = 4). (**E**– **H**) Experiments with cell type-specific antibody treatment of either collagen-adherent platelets or RBCs with: an anti-CD36 mAb (**E**, platelets: 2 µg/mL; **F**, RBCs: 2 µg/mL), an anti-TSP-1 mAb (**G**, platelets: 2 µg/mL), abciximab (**H**, 10 µg/2 million cells) and IgG-treated control (*n* = 3–5). Data are represented as mean values ± SEM. \**P* < 0.05; \*\**P* < 0.01 tested by student’s t-test (**C, E**–**H**) and one-way ANOVA with Sidak’s multiple comparison (**D**). Plts, platelets; RBCs, red blood cells.

Next, the role of CD36 for RBC recruitment was investigated. The number of transient adherent RBCs showed a significant reduction when either platelets or RBCs were treated with a blocking anti-CD36 antibody (Figure 4E and F, Supplemental Videos 1–4). Inhibition of TSP-1 resulted in a significant reduction of transient adherent RBCs (Figure 4G, Supplemental Videos 5 and 6). In contrast, blocking of integrin α_IIb_β_3_ using abciximab did not alter the recruitment of RBCs to collagen-adherent platelets, suggesting that this integrin does not participate in the recruitment of RBCs (Figure 4H). The inhibition of either FasL by hDcR3 or FasR by antibody treatment led to a significantly reduced number of transient adherent RBCs compared to IgG-treated controls (Supplemental Figure 5A and B, Supplemental Videos 7–10). Taken together, the interaction of RBCs and platelets via CD36–TSP-1 and FasL–FasR are responsible not only for the increase externalization of PS on the membrane of RBCs and platelets but also for the active recruitment of RBCs to collagen-adherent platelets under flow conditions. Of note, integrin α_IIb_β_3_ does not appear to play a role in the recruitment of RBCs.

### CD36–TSP-1-mediated interaction of platelets and RBCs is important for platelet activation and thrombus formation *in vivo*

To investigate whether the CD36–TSP-1-mediated interaction of platelets and RBCs plays a role in shear-dependent adhesion of platelets at sites of vascular injury, platelet recruitment to the ligated carotid artery was analyzed by *in vivo* fluorescence microscopy. For visualization, platelets from donor mice were fluorescently labeled and quantified. As shown in Figure 5A, significantly fewer platelets from CD36-deficient mice stably adhered to the ligated carotid artery at 20 min post-ligation (Figure 5A, Supplemental Videos 11 and 12). Reduced stable adhesion of TSP-1-deficient platelets was also observed following vessel injury by ligation (Figure 5B, Supplemental Videos 13 and 14), suggesting that both CD36 and TSP-1 play an important role in the recruitment of platelets to injured vessel walls. Next, we investigated the CD36–TSP-1 axis in shear-dependent arterial thrombus formation *in vivo*, as shown recently for FasR and FasL (21). To distinguish between the roles of platelet CD36 and erythroid CD36, we used conditional knock-out mice with a CD36 deficiency restricted to platelets (*Cd36^fl/fl^-Pf4-Cre*) or RBCs (*Cd36^fl/fl^-Hbb-Cre*). Topical application of 20% FeCl_3_ was applied to exposed mesenteric arterioles to induce vessel injury and arterial thrombus formation, leading to full occlusion of the injured vessel in control mice (*Cd36^fl/fl^-Pf4-Cre*^−^) but not in *Cd36^fl/fl^-Pf4-Cre^+^* mice (Figure 5C–E, Supplemental Videos 15 and 16), confirming published data regarding the importance of platelet CD36 in thrombus formation under conditions of high-shear (8). However, thus far, the role of CD36 in thrombus formation has been assigned to platelets alone. Therefore, we analyzed the role of erythroid CD36 in thrombus formation under high-shear conditions as well. As shown in Figure 5F–H, mice with a CD36 deficiency restricted to RBCs (*Cd36^fl/fl^-Hbb-Cre*) showed delayed initiation of thrombus formation (by trend, Figure 5G) and no occlusion of the vessel at the end of the 40-min observation period (Figure 5F and H, Supplemental Videos 17 and 18). These results clearly indicate a role for the CD36–TSP-1 axis, and specifically erythroid CD36 in arterial thrombus formation *in vivo*.

**Figure 5.**
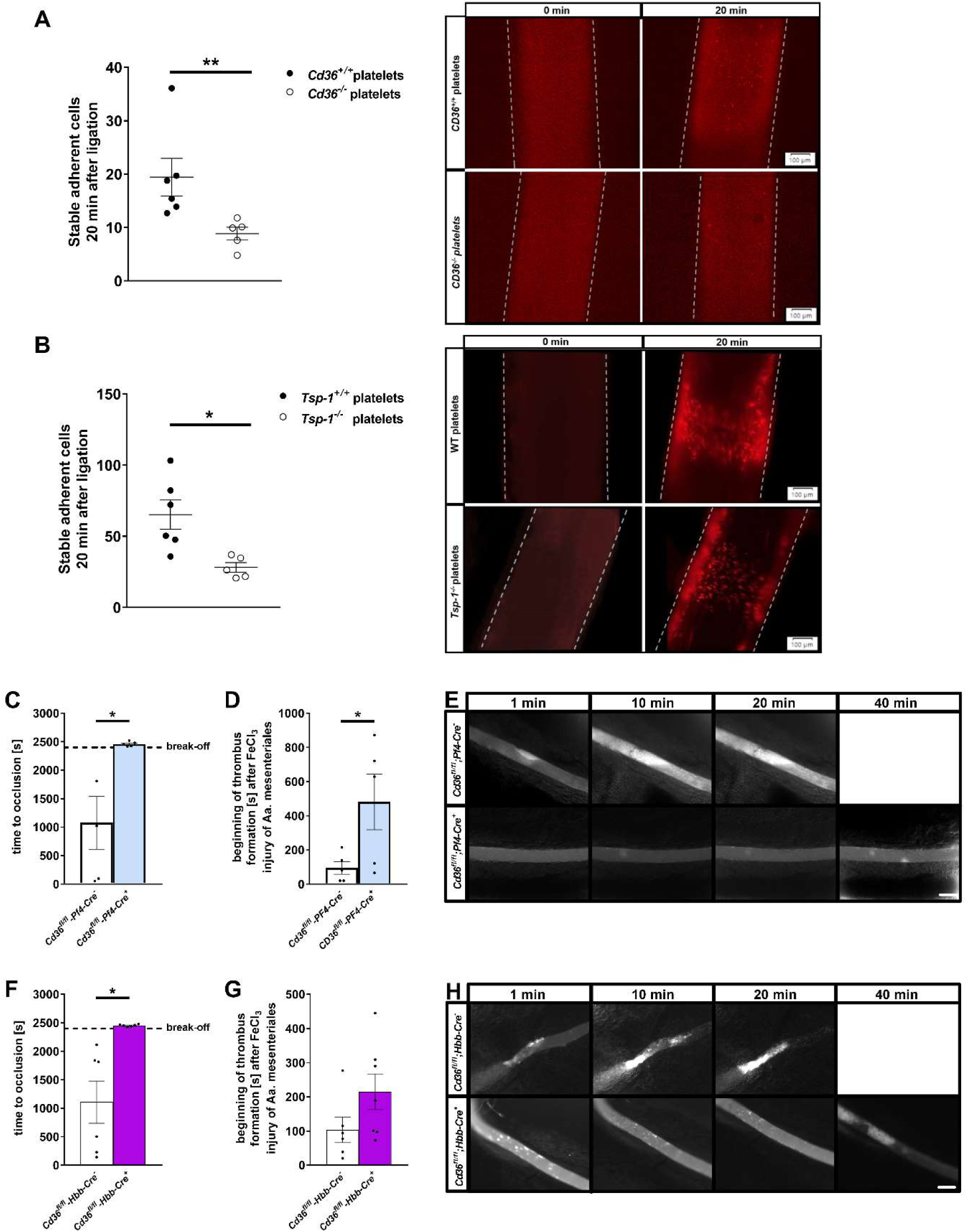
Deletion of TSP-1 and CD36 reduces platelet adhesion *in vivo*. (**A** and **B**) Quantification and images of stable adherent platelets 20 min after carotid artery ligation *in vivo*. (**A**) Platelets from *Cd36*^−/–^ and *Cd36*^+/+^ donor mice were stained and intravenous injected to recipient WT mice. Vessel injury was induced in WT (littermates, *Cd36*^+/+^) mice (*n* = 5–6) (**B**) Platelets from *Tsp-1*^−/−^ and *Tsp-1*^+/+^ donor mice were stained and intravenous injected to recipient WT mice. Vessel injury was induced in WT (C57BL/6J, *Tsp-1*^+/+^) mice (*n* = 5–6). Images represent carotid artery vessel wall, which is outlined using dotted lines (**A** and **B**). (**C**– **H**) FeCl_3_-induced injury of mesenteric arterioles in *Cd36^fl/fl^-Pf4-Cre* and *Cd36^fl/fl^-Hbb-Cre mice* (*n* = 5–7). Percentage of mice with or without occlusion after 40 min (**C** and **F**), beginning of thrombus formation (**D** and **G**) and representative intravital microscopy images at defined time points (**E** and **H**). Scale bar: 100 μm. Data are represented as mean values ± SEM. \**P* < 0.05 tested by Mann-Whitney test.

### Enhanced procoagulant activity of RBCs and platelets in patients with abdominal aortic aneurysm

To date, little is known regarding the impact of platelet–RBC interactions on arterial thrombosis in cardiovascular diseases. Recently, we have provided evidence for the direct contact of platelets and RBCs in murine and human thrombi, as seen in surgical specimens of patients after thrombectomy (21). To further investigate the impact of platelet–RBC interactions and the procoagulant activity of these cells, we analyzed platelet–RBC interactions and procoagulant activity in an experimental mouse model of AAA (the pancreatic porcine elastase [PPE] model) and in patients with AAA. We recently detected elevated platelet activation and platelet-mediated inflammation in mice and patients with AAA (26). Here, histological analysis at day 28 after elastase infusion revealed that platelets and RBCs had migrated into the abdominal aorta of PPE mice (Figure 6A). At day 28, most of the cells were detected in the adventitia (Figure 6A). Furthermore, we detected enhanced procoagulant activity of RBCs in the presence of CRP-activated platelets (Figure 6B) and platelets under resting and activated conditions (Figure 6C). In patients with AAA, enhanced coagulation that correlated with the size of the AAA was detected (27). To evaluate the pathophysiologic relevance of platelet– RBC interactions in these processes, we characterized platelets and RBCs by flow cytometry using whole blood from patients with AAA and age-matched healthy volunteers and observed increased procoagulant activity of RBCs (Figure 6D) and platelets (Figure 6E) under resting conditions. Procoagulant activity of platelets was also enhanced by low doses of CRP, while high concentrations of CRP did not induce alterations in annexin V binding, suggesting that the threshold to expose PS at the platelet and RBC membranes was reduced in these patients (Figure 6D and E). To determine activation-dependent upregulation of integrin α_IIb_β_3_ at the platelet surface, we measured CD61 on the membrane of resting and activated platelets (Figure 6F). Low, intermediate, and high concentrations of ADP, CRP, or PAR4 peptide increased amount of integrin β_3_ at the platelet surface in a dose-dependent manner. In addition, enhanced integrin exposure under resting conditions and with high concentrations of ADP and CRP was observed in AAA patients compared to age-matched controls (Figure 6F). As shown in Figure 6G, we detected reduced platelet–RBC aggregates in AAA patients that may have been due to reduced platelet and RBC counts in these patients. (Supplemental Figure 6A and B). Because most AAA patients develop an intraluminal thrombus (ILT) (28, 29), reduced blood cell counts and aggregates may be due to elevated consumption of platelets and RBCs at the interface of blood flow and the ILT in the aorta.

**Figure 6.**
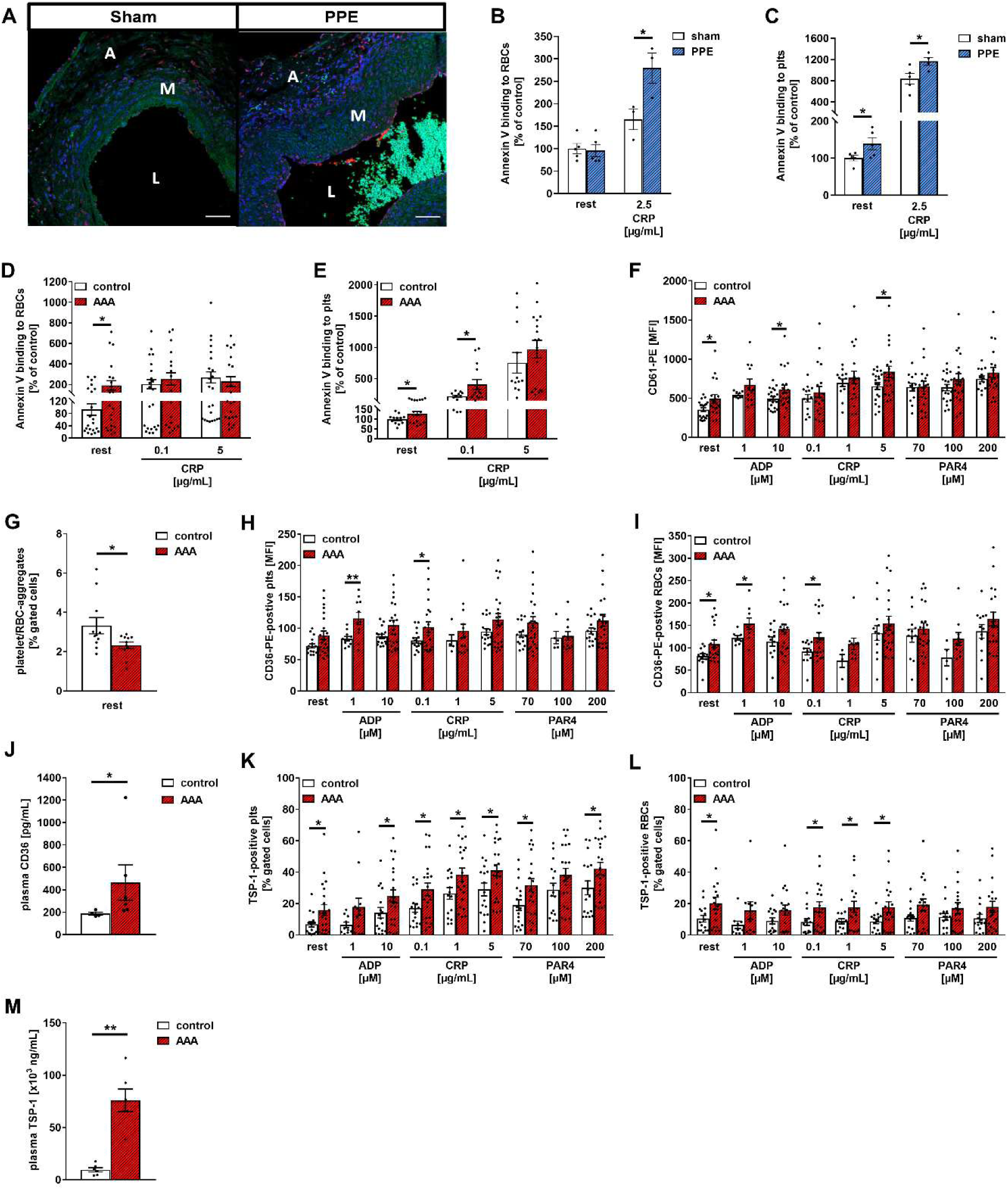
Enhanced procoagulant surface of RBCs and platelets in PPE-operated mice and AAA patients. (**A**) Representative immunofluorescence (IF) images of platelets (CD42b/anti-GPIbα, red), nuclei (DAPI, blue), elastin (autofluorescence, green) and RBCs (autofluorescence, green) in aortic tissue sections of PPE-operated mice (sham or PPE) at day 28 after surgery (*n* = 3). (**B** and **C**) Annexin V binding to RBCs (**B**) or platelets (**C**) from sham- or PPE-operated mice at day 28 under resting and CRP-stimulated conditions determined by flow cytometry (*n* = 4–5), indicated as percent-gated cells normalized to controls. (**A**–**M**) Human whole blood was collected from AAA patients and age-matched controls, stimulated with indicated agonists and determined by flow cytometry. Concentrations of FasL, CD36 and TSP-1 were determined in plasma samples by ELISA. Age-matched controls (≥ 60 years old) were used as control group. (**D** and **E**) Annexin V binding to RBCs (**A**) and platelets (**B**) under resting and CRP-stimulated conditions (controls, *n* = 12–23; AAA, *n* = 12–20), indicated as percent-gated cells normalized to controls. (**F**) Externalization of the integrin β_3_ subunit (CD61-PE) on the surface of platelets (controls, *n* = 13–19; AAAs, *n* = 11). (**G**) Aggregate formation of platelets (CD42^+^) and RBCs (CD235^+^) in whole blood (controls, *n* = 10; AAAs, *n* = 11). (**H** and **I**) CD36 expression on the surface of platelets (**H**) and RBCs (**I**) (controls, *n* = 5–16; AAAs, *n* = 11–23). (**J**) Analysis of soluble CD36 in plasma samples of AAA patients and respective controls (controls, *n* = 5; AAAs, *n* = 6). (**K and L**) TSP-1 binding to platelets (**K**) and RBCs (**L**) (controls, *n* = 10–17; AAAs *n* = 12–22). (**M**) Analysis of soluble TSP-1 in plasma samples of AAA patients and respective controls (*n* = 6). Data are represented as mean values ± SEM. \**P* < 0.05; \*\**P* < 0.01 tested by multiple t-test (**B**–**F, H, I, K, L**) and Mann-Whitney test (**G, J, M**). Plts, platelets; RBC, red blood cells; A, adventitia; L, lumen; AAA, abdominal aortic aneurysm; control, age-matched controls; PPE, porcine pancreatic elastase (infusion); ADP, adenosine diphosphate; CRP, collagen related peptide; PAR4, protease-activated receptor 4 activation peptide; MFI, mean fluorescence intensity.

To investigate whether or not the CD36- and TSP-1-mediated interaction of platelets and RBCs is altered in AAA, we determined the exposure of CD36 on the membrane of platelets (Figure 6H) and RBCs (Figure 6I) by flow cytometry. Differences in the amount of CD36 on the membrane of these cells with low concentrations of ADP or CRP as well as under resting conditions of RBCs were detected between AAA patients and age-matched healthy volunteers (Figure 6H and I). However, stimulation by PAR4 peptide revealed no alterations of CD36 externalization on the membrane of platelets and RBCs. In addition, plasma levels of soluble CD36 were enhanced in AAA patients compared to age-matched controls, suggesting enhanced activity of the receptor (Figure 6J). Compared to age-matched controls, we detected enhanced binding of TSP-1 to platelets and RBCs from AAA patients under resting conditions and after stimulation of platelets with ADP, CRP, or PAR4 peptide (Figure 6K and L). In line with enhanced levels of soluble CD36, we detected elevated levels of TSP-1 in the plasma of AAA patients compared to controls (Figure 6M), suggesting that the activation of both CD36 (platelet-derived and erythroid) and TSP-1 is elevated in AAA.

We also quantified the amount of FasR exposed on the membrane of RBCs in the presence of resting or stimulated platelets using flow cytometry (Supplemental Figure 6C). Neither the stimulation of platelets with different concentrations of CRP, PAR4 peptide, or ADP nor the presence of resting platelets altered the amount of FasR on the membrane of RBCs from patients with AAA compared to age-matched healthy controls (Supplemental Figure 6C), suggesting that the exposure of FasR is not regulated by interaction with platelets or by AAA pathology. In contrast to FasR exposure, the presence of FasL at the platelet surface is activation-dependent (21, 30). In AAA patients, we detected enhanced exposure of FasL on the membrane of resting platelets compared to age-matched controls but unaltered FasL levels on the platelet membrane upon platelet activation using indicated agonists (Supplemental Figure 6D). However, the plasma levels of soluble FasL were unaltered in AAA patients compared to age-matched controls (Supplemental Figure 6E). Taken together, enhanced exposure of platelet FasL, increased exposure of CD36 on platelets and RBCs, and increased binding of TSP-1 to both cell types implies elevated interactions of platelets and RBCs in AAA that may be responsible for enhanced procoagulant activity of RBCs (Figure 6D) and platelets (Figure 6E).

### Deposition of platelets, RBCs and TSP-1 in the aortic wall and in the intraluminal thrombus of AAA patients

We next analyzed whether enhanced procoagulant activity of RBCs and platelets (Figure 6D and E) is accompanied by the migration of platelets and RBCs into the aortic wall and the ILT of AAA patients, as previously observed in experimental mice with AAA (PPE mice, Figure 6A). In Figure 7A, we provide evidence for the presence of RBCs, platelets, and TSP-1 in AAA specimens. Platelets and RBCs were detected in the aortic wall of patients with AAA (Figure 7A, upper right panel and Supplemental Figure 7B, upper middle panel). Furthermore, TSP-1 had accumulated in different regions of the aortic wall, including the tunica intima, the tunica media, and the tunica adventitia (Figure 7A, lower right panel, Supplemental Figure 7C, upper middle panel). Accumulation of TSP-1 was also found in the ILT of AAA patients (Figure 7B, upper right panel, Supplemental Figure 7C, upper middle panel). In addition, high numbers of platelets and RBCs were detected in the ILT of these patients (Figure 7B, upper left panel, Supplemental Figure 7B, middle panel). TSP-1 had also accumulated in thrombi that were isolated from patients after thrombectomy (Figure 7B, lower right panel, Supplemental Figure 7C, lower middle panel). However, accumulation of TSP-1 in these thrombi was relatively low compared to the massive accumulation of TSP-1 in the ILT of AAA patients. Furthermore, platelets and RBCs were detected in the thrombi of patient specimens after thrombectomy (Figure 7B, lower left panel, Supplemental Figure 7B, lower middle panel) as shown previously earlier by HE staining and electron microscopy (21). Taken together, these data imply that platelets and RBCs as well as TSP-1 accumulate in the aortic wall of AAA patients. However, how these processes contribute to aneurysm formation or progression remains elusive.

**Figure 7.**
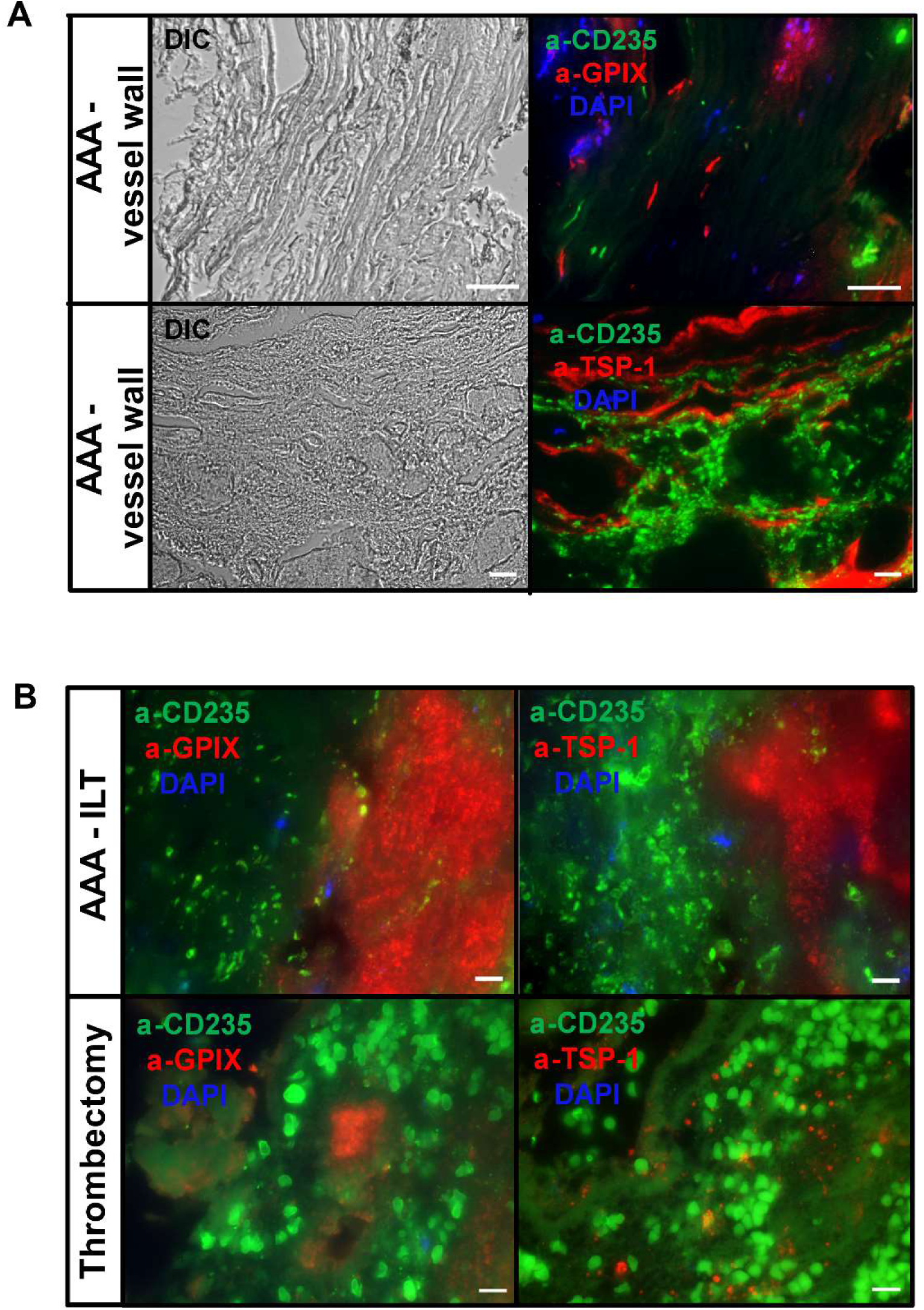
Deposition of platelets, RBCs and TSP-1 in the aortic wall and the intraluminal thrombus of AAA patients. (**A** and **B**) Paraffin-embedded sections (5 µm sections) of the aortic wall (**A**) and the ILT (**B**) (upper panels) from AAA patients, and of thrombi isolated from patients who underwent thrombectomy (**B**) (peripheral thrombi, lower panel) were stained with an anti-GPIX mAb (20 µg/mL) to detect platelets, an anti-CD235 mAb (0.5 µg/mL) to detect RBCs, and an anti-TSP-1 mAb (20 µg/mL) to detect accumulated TSP-1 (*n* = 3–4). Additionally, fluorescence emission at 488 nm showed autofluorescence elastic lamina in samples of the vessel wall. 4’, 6-diamidino-2-phenylindole (DAPI) was used to stain nuclei. Representative DIC and fluorescence images with 400-fold (**A**, upper and middle panels, scale bar: 50 µm) and 1000-fold (**A,** lower panel; **B**, all images, scale bar: 10 µm) magnification are shown. AAA, abdominal aortic aneurysm; ILT, intra luminal thrombus.

### Genetic deletion of TSP-1 reduces AAA progression through platelet activation, procoagulant surface, and the recruitment of inflammatory cells

To further analyze the role of TSP-1 in AAA, we induced experimental AAA in TSP-1-deficient mice utilizing the PPE model. We found that TSP-1 deficiency attenuated the diameter progression of AAA over a 28-day time course by ultrasound tracking (Figure 8A). Percentage survival of mice was unaltered between groups (Figure 8B), but TSP-1-deficient mice showed reduced AAA incidence (Figure 8C). Next, we determined the procoagulant activity of both platelets and RBCs. The procoagulant activity of RBCs was enhanced in naive TSP-1-deficient mice but unaltered in PPE mice at day 28, suggesting reduced procoagulant activity of RBCs in experimental AAA compared to naive controls (Figure 8D). We detected unaltered PS exposure of platelets in naive mice but reduced annexin V binding to TSP-1-deficient platelets at day 28 post-surgery (Figure 8E). Furthermore, we detected reduced numbers of platelets and RBCs in the aortic wall of TSP-1-deficient mice at day 28 post-surgery (Figure 8F). This was accompanied by reduced integrin α_IIb_β_3_ exposure at the surface of TSP-1-deficient platelets under resting and stimulated conditions (Figure 8G), reduced integrin α_IIb_β_3_ activation of platelets in response to various concentrations of CRP and PAR4 peptide determined by flow cytometry (Figure 8H). In addition, P-selectin exposure was reduced in response to low and intermediate concentrations of PAR4 peptide using TSP-1-deficient platelets in washed whole blood from PPE mice (Figure 8 H). However, the number of platelet–RBC aggregates in the plasma was unaltered in TSP-1-deficient PPE mice (Figure 8I). In contrast, we detected a reduced number of platelet–leukocyte (Figure 8J, left panel) and platelet–neutrophil aggregates (Figure 8J, right panel) in TSP-1-deficient PPE mice, suggesting that TSP-1 is important for platelet-induced activation of leukocytes in AAA, in line with the role of TSP-1 in the inflammatory response in PPE mice (31). However, this study did not adress the role of platelets. Taken together, our data have revealed an important impact of TSP-1 on procoagulant activity of platelets and RBCs as well as on the upregulation and activation of integrin α_IIb_β_3_ in AAA.

**Figure 8.**
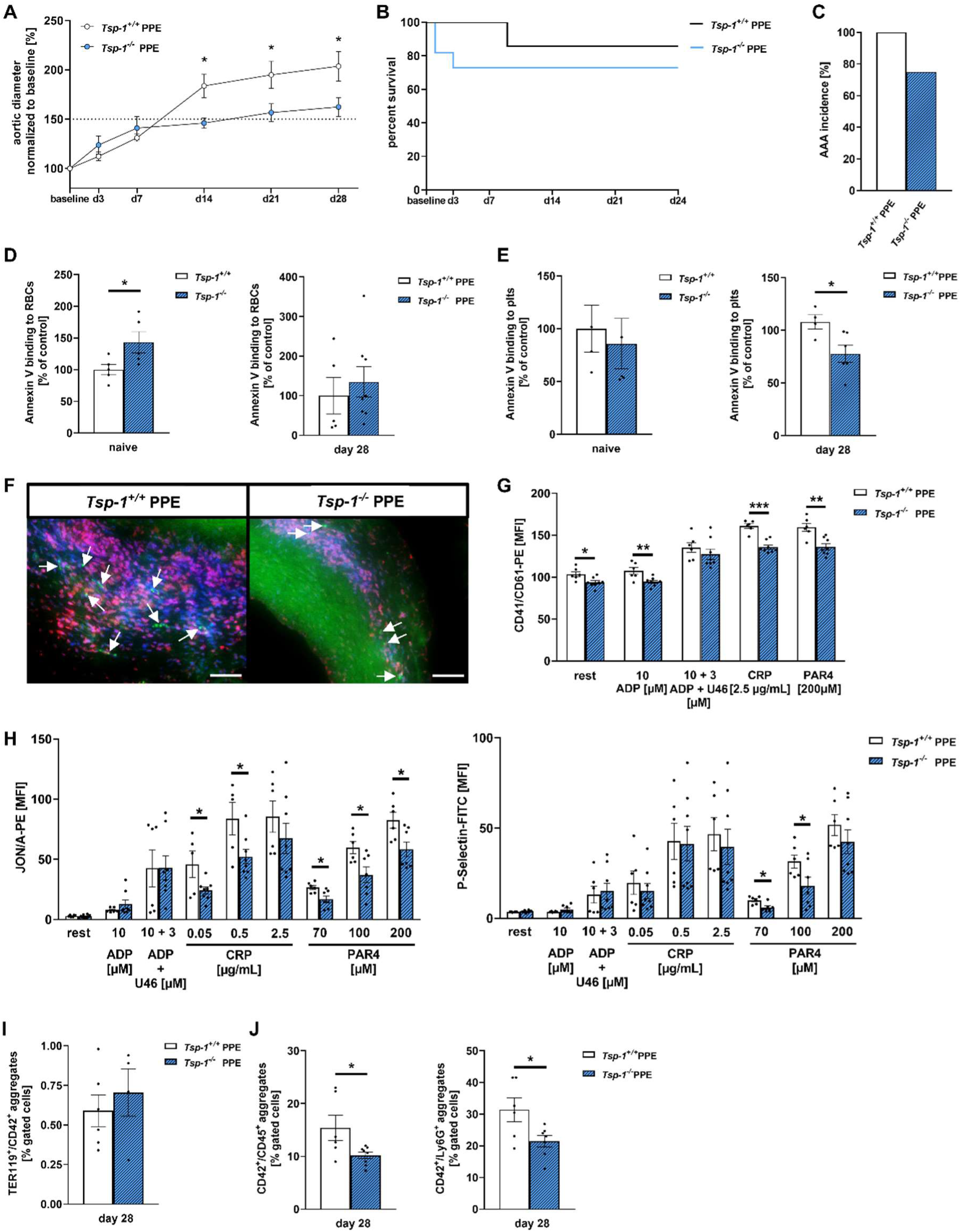
Genetic deletion of TSP-1 reduces AAA progression through platelet activation, procoagulant surface and the aggregate formation with inflammatory cells. (**A**) Ultrasound measurements were performed prior surgery (baseline) and at days 3, 7, 14, 21 and 28 to monitor aneurysm diameter progression (controls, *n* = 6–7; PPE-operated mice, *n* = 7–12). (**B**) Survival rate of *Tsp-1^+/+^* mice over 28 days compared to *Tsp-1*^−/–^ mice. (**C**) Incidence of AAA formation of *Tsp-1^+/+^* mice compared to *Tsp-1*^−/–^ mice (AAA diameter ≥ 150%) at day 28 post-surgery. (**D** and **E**) Annexin V binding to RBCs (**D**) and platelets (**E**) in whole blood from naive or *Tsp-1*^−/–^ PPE-operated mice at day 28, determined by flow cytometry (*n* = 4–8). Data are represented as percent-gated cells normalized to controls. *Tsp-1^+/+^*mice served as controls. (**F**) Representative immunofluorescence (IF) images of platelets (CD42b/anti-GPIbα, red), nuclei (DAPI, blue), elastin (autofluorescence, green) and RBCs (autofluorescence, green) in aortic tissue sections of PPE-operated mice *Tsp-1^+/+^*, respectively *Tsp-1*^−/–^ mice at day 28 post-surgery (*n* = 4). (**G**) Upregulation of integrin α_IIb_β_3_ (CD41/CD61-PE) (*n* = 6–8). (**H**) Active integrin α_IIb_β_3_ externalization (JON/A-PE, left) and degranulation marker (P-selectin-FITC, right) of *Tsp-1^+/+^*mice compared to *Tsp-1*^−/–^ mice at day 28 with indicated agonists, determined by flow cytometry (*n* = 5–8). (**I**) Aggregate formation of platelets (CD42^+^) with RBCs (TER119^+^) in whole blood samples (*n* = 5–6). (**J**) Aggregate formation of platelets (CD42^+^) with leucocytes (CD45^+^, left panel), respectively neutrophils (Ly6G^+^, right panel) in whole blood samples of PPE-operated *Tsp-1^+/+^* and *Tsp-1*^−/–^ mice at day 28, determined by flow cytometry (*n* = 6–8). Data are represented as mean values ± SEM. \**P* < 0.05, \*\**P* < 0.01, \*\*\**P* < 0.001 tested by two-way ANOVA with Sidak’s multiple comparison test (**A**), unpaired student‘s t-test or multiple t-test (**D**–**J**). Plts, platelets; RBC, red blood cells; A, adventitia; M, media; L, lumen; PPE, porcine pancreatic elastase (infusion); MFI, mean fluorescence intensity; ADP, adenosine diphosphate; CRP, collagen-related peptide; PAR4, protease-activated receptor 4 activating peptide.

## DISCUSSION

In this study, we have shown that CD36 on RBCs supports thrombus formation on immobilized collagen and on collagen–TSP-1. The impact of erythroid CD36 in thrombus formation is important under high-shear conditions and may reflect the ability of TSP-1 to bind to GPIb and to RBCs. Under static conditions, CD36 on RBCs and platelets is important for the procoagulant activity of RBCs; however, under conditions of arterial shear, only erythroid CD36 plays a role in thrombus generation by RBCs but not of platelets. TSP-1 binding to platelets and RBCs is highly increased under flow conditions and mediated at least in part by erythroid CD36. Integrin α_IIb_β_3_ was identified as another ligand for FasR on RBCs to promote procoagulant activity. Platelet–RBC interactions via FasL–FasR and CD36–TSP-1 not only promote procoagulant activity of RBCs but also enhance platelet aggregation and the recruitment of RBCs to the growing thrombus. Thus, CD36 and TSP-1 are indispensable for the recruitment of platelets to the arterial wall.

We have also provided evidence that erythroid CD36 plays an essential role in occlusive thrombus formation, as CD36 deficiency restricted to RBCs resulted in almost no occlusion of injured mesenteric arterioles as compared to CD36 deficiency restricted to platelets. Enhanced procoagulant activity and the recruitment of RBCs and platelets to the aortic wall in experimental mice and to the ILT in patients with AAA suggest that platelet–RBC interactions via CD36–TSP-1 are crucial for the progression of AAA. In experimental PPE mice, TSP-1 is important in platelet–RBC interactions and procoagulant activity of these cells, as TSP-1-deficient mice showed reduced aortic aneurysm progression, reduced platelet activation, and less procoagulant activity of platelets and RBCs.

In the absence and in the presence of hyperlipidemia, CD36 plays an important role in thrombosis (32–34). Specifically, CD36 on platelets promotes platelet adhesion to TSP-1 and platelet activation, including integrin α_IIb_β_3_ activation and P-selectin exposure (9). Furthermore, thrombus formation on a collagen–TSP-1 matrix has been shown to be dependent on CD36, as TSP-1 represents a ligand for CD36 on platelets (8, 9). In this study, we have confirmed enhanced thrombus formation on a collagen–TSP-1 matrix compared to collagen alone using PRP and RBCs but not using whole blood, as done in earlier studies. In line with previous data, we detected reduced thrombus formation on collagen–TSP-1 when we blocked CD36 on either platelets or RBCs, suggesting that thrombus formation on collagen–TSP-1 is mediated by CD36 on both platelets and RBCs (Figure 2A and B) (8, 9).

Thus, a role for CD36 in thrombus formation on collagen alone has not been detected, either by using an anti-CD36 blocking antibody or in analyzing CD36-deficient whole blood (8, 9). In the present study, we found reduced thrombus formation on collagen under flow. In contrast to other published work (8, 9), we used a shear rate of 1,700 s^−1^. Under these conditions, we observed reduced thrombus formation on collagen when we applied an anti-CD36 blocking antibody to whole blood or blocked CD36 on RBCs, while no differences in thrombus formation were observed after blocking platelet CD36 (Figure 1D). Thus, the differences from previously published data are due to the shear rates used in this study (Supplemental Figure 1A and B). Whereas the group of Heemskerk perfused human or murine whole blood over a collagen matrix with a shear rate of 1,000 s^−1^, we used a shear rate of 1,700 s^−1^, at which thrombus formation at least partially depend on GPIb (Figure 1D) (9). However, the impact of CD36 in thrombus formation on collagen was not related to platelet CD36 but rather to erythroid CD36 expression, as the blocking of CD36 only on RBCs reduced surface coverage on collagen (Figure 1D, 2A and B). Thus, the impact of RBC CD36 in thrombus formation on collagen depends on GPIb-mediated platelet adhesion under high-shear conditions. In contrast, CD36-mediated thrombus formation on collagen–TSP-1 is also relevant at a shear rate of 1,000 s^−1^ (8). This might be due to the fact that TSP-1–GPIb binding is an alternative mechanism to vWF binding to GPIb, which is important for platelet adhesion under pathophysiological high-shear rates (35). In addition, TSP-1 released from platelets modulates thrombosis in the presence of vWF (36). Thus, the vWF–GPIb axis might be involved in platelet–RBC interactions via CD36–TSP-1 and RBCs seem to be essential for stable thrombus formation under high-shear via TSP-1 binding.

Furthermore, our data provide evidence for binding of platelet-released TSP-1 to erythroid CD36 to modulate procoagulant activity of RBCs, which is important to promote thrombus formation on collagen. TSP-1 binding to platelets and RBCs isolated from the thrombus was dramatically enhanced compared to the flow-through using either human or murine blood (Figure 2F and L). Congruently, our data indicate that CD36 plays a role in the binding of TSP-1 to RBCs inside the thrombus but not in the flow-through (Figure 2L), for procoagulant activity of RBCs (Figure 1E), and in thrombus formation (Figure 1D). Because CD36 on RBCs is only important for thrombus formation on collagen under high-shear, it is plausible that TSP-1 binding by erythroid CD36 is modulated by shear as well.

Nergiz-Unal and colleagues have shown that platelet CD36 is involved in human platelet activation but not in PS exposure upon static adhesion to TSP-1. In contrast, CD36 has been shown to play a role in the exposure of PS upon platelet adhesion and thrombus formation on a collagen–TSP-1 matrix under flow conditions (shear rate 1,000 s^−1^). However, annexin V binding to the thrombus was measured by including the exposure of PS of RBCs and platelets (8, 9). In contrast, annexin V binding to the growing thrombus was not altered when whole blood was perfused over collagen alone. Here, we also detected no alterations in platelet PS exposure upon platelet adhesion to collagen when CD36 on platelets had been blocked by antibody treatment, under either static or dynamic conditions (Figure 1B and F). These results were confirmed using CD36-deficient platelets from transgenic mice (Figure 1J and L). However, when the blockage of CD36 was restricted to RBCs, we found reduced PS exposure of human platelets under static (Figure 1B) but not under dynamic conditions (Figure 1F). In contrast, PS exposure of RBCs was reduced when CD36 was either blocked or deleted on platelets or RBCs under static conditions (Figure 1C and I). Under flow conditions, blocking or genetic deletion of erythroid CD36 resulted in reduced exposure of PS of RBCs (Figure 1E and K). Thus, the exposure of RBCs and platelets to shear may be important for the procoagulant activity of RBCs mediated by CD36. Because the exposure of PS strongly depends on shear conditions (37), we assume that mainly erythroid CD36 is important for the procoagulant activity of RBCs. This conclusion is supported by the results shown here, demonstrating that TSP-1 binding to erythroid CD36 is also triggered by high-shear.

Our results provide evidence of a role for integrin α_IIb_β_3_ in the interaction of platelets and RBCs, suggesting two independent mechanisms for platelet–RBC interactions to induce procoagulant activity of RBCs via activation of erythroid FasR. Integrin α_IIb_β_3_ has previously been shown to interact with erythroid ICAM-4, which is important for the incorporation of RBCs into a thrombus (38, 39). Here, we have shown that integrin α_IIb_β_3_ is important not only for the exposure of PS at the platelet surface (37) but also for the procoagulant activity of RBCs. Of note, our data suggest a synergistic effect of FasL and integrin α_IIb_β_3_ in inducing the activation of FasR at the RBC membrane, as PS exposure of murine FasL-deficient platelets and RBCs was further reduced in the presence of an antibody blocking integrin α_IIb_β_3_ (Supplemental Figure 3J and K). The upregulation of integrin α_IIb_β_3_ exposure at the platelet surface shown here in the presence of RBCs depends on erythroid receptors that play a role in platelet–RBC interactions, such as CD36 and FasR (Figure 3B, Supplemental Figure 3A and B), but not on platelet-derived ligands, such as TSP-1 or FasL (Figure 3C, Supplemental Figure 3C and D). In addition to the role of integrin α_IIb_β_3_ in PS exposure, we have also provided evidence that RBCs support platelet aggregation. This support may reflect the ability of RBCs to induce the upregulation of integrin α_IIb_β_3_ at the platelet surface, which may enhance fibrinogen binding of platelets. Moreover, integrin α_IIb_β_3_ modulates procoagulant activity of not only platelets but also of RBCs (Supplemental Figure 3I). Thus, enhanced thrombin generation in the presence of RBCs may support platelet aggregation.

A role for integrin α_IIb_β_3_ in the recruitment of RBCs to the growing thrombus has been described previously (38). In this study, we found that human integrin α_IIb_β_3_ had no impact on the recruitment of RBCs to collagen-adherent platelets (Figure 4). However, the exposure of PS of both platelets and RBCs plays a crucial role in the recruitment of RBCs under arterial shear. Thus, it is not surprising that the receptor–ligand pairs that are involved in platelet–RBC interactions, such as FasL–FasR and CD36–TSP-1 are involved in the recruitment of RBCs to the growing thrombus. Consequently, CD36 deficiency restricted to RBCs resulted in almost no occlusion of mesenteric arterioles, as previously observed for platelet CD36 (Figure 5).

We have recently provided evidence for a role of platelet–RBC interactions via FasL–FasR in pathological thrombus formation (21). The data presented here have revealed that platelet– RBC interactions are also important modulators in the formation and progression of AAA. Platelets and RBCs from experimental PPE mice and AAA patients showed enhanced procoagulant activity (Figure 6) that may reflect the enhanced plasma thrombin levels that have recently been described in AAA patients (40, 41). Furthermore, we detected unaltered soluble FasL but enhanced plasma levels of TSP-1 and CD36 in AAA patients, suggesting elevated activity of the CD36–TSP-1 axis in AAA pathology. In addition, enhanced FasL externalization, increased CD36 and TSP-1 binding to resting and mildly or strongly activated platelets from AAA patients suggests that platelet–RBC interactions may be enhanced, as evidenced by enhanced procoagulant activity of RBCs and platelets and the presence of these cells in the aortic wall and in the ILT of AAA patients (Figure 6 and 7). Although a role for TSP-1 in the inflammatory response in AAA has been shown previously (31), we have demonstrated here a crucial role for TSP-1 in procoagulant activity of platelets and RBCs and in the upregulation and activation of integrin α_IIb_β_3_ in AAA, strengthened by the analysis of TSP-1-deficient mice in experimental AAA (Figure 8).

To gain further insight into the pathology of AAA, a genomic approach was undertaken, finding TSP-1 and the subunit α_IIb_ to be highly upregulated in AAA (42). A contribution of procoagulant activity of RBCs under pathological conditions has previously been shown in patients with sickle-cell disease and nephrotic syndrome (43, 44). Of note, CD36 plays a role in the recruitment of PS-positive microparticles into the growing thrombus (32) and has been associated with inflammatory and pro-thrombotic properties in acute coronary syndrome (45). Thus, it is not surprising that we could provide evidence for a contribution of procoagulant platelets and RBCs, integrin α_IIb_β_3_, CD36, and TSP-1 to thrombus formation and AAA progression in mice and humans.

Taken together, our results provide strong evidence of a mechanism by which platelets and RBCs interact in thrombus formation in arterial thrombosis and AAA pathology. As shown previously for platelet–RBC interaction via FasL–FasR, CD36–TSP-1-mediated cell–cell interaction results not only in PS exposure and procoagulant activity of RBCs, thus contributing to the active role of RBCs in thrombus formation, but also in the active recruitment of RBCs to collagen-adherent platelets and to platelet aggregation under static conditions. Moreover, we have identified integrin α_IIb_β_3_ as another ligand in addition to FasL for erythroid FasR; this observation may explain the recent description of moderate effects of FasL in contrast to FasR in platelet–RBC interactions (21). In an initial translational approach, we have provided evidence for a crucial role of FasL–FasR- and CD36–TSP-1-mediated platelet–RBC interactions in AAA, as enhanced procoagulant activity of platelets and RBCs was detected in AAA patients with enhanced exposure of FasL and TSP-1 at the platelet surface. Because no clinical therapy for AAA patients is currently available, we believe that interfering with the interaction of platelets and RBCs – specifically with the CD36–TSP-1 or FasL–FasR axis – may be an innovative and promising therapeutic approach to protect AAA patients from surgery or aneurysm rupture. In future, our investigations may help to establish a new plasmatic coagulation assay or diagnostic tool to determine risk stratification in AAA progression.

## METHODS

### Study approval

All animal experiments were conducted according to the Declaration of Helsinki and German law for the welfare of animals. The protocol was reviewed and approved by the local government authority. Heinrich-Heine-University Animal Care Committee and by the State Agency for Nature, Environment and Consumer Protection of North Rhine-Westphalia (LANUV), Recklinghausen, NRW, Germany; Permit Numbers: 81-02.04.2018.A409, 81-02.04.2020.A284, 84-02.05.40.16.073, and 81-02.05.40.21.041 and by the *Landesuntersuchungsamt* Rhineland-Pfalz, Koblenz, Germany; Permit number: G16-1-013. Animals were housed in a temperature and humidity-controlled room under a 12-h light/dark cycle (6:30 am–6:30 pm). Experiments using human tissues and blood were reviewed and approved by the Ethics Committee of the Heinrich-Heine-University, who approved the collection and analysis of the tissue. Permitted ethical votes: 2018-140-KFogU, 5731R (2018-222_1) and 2018-248-FmB. Subjects provided informed consent prior to their participation in the study (patient consent). The study was conducted in accordance with the principles of the Declaration of Helsinki and the International Council for Harmonization Guidelines on Good Clinical Practice.

### Animals

Specific pathogen-free C57BL/6J mice were obtained from Janvier Labs. Gene-targeted mice lacking Fas receptor (*Fas*^−/–^) were obtained from The Jackson Laboratory (C57BL/6J.MRL-FAS^lpr^). *Cd36*^−/–^ mice were obtained from Dr. Hadi-Al Hasani (German Diabetes Center) and originally from Maria Febbraio. Fasl^fl/fl^ mice were crossed to Pf4-Cre mice, which had been purchased from The Jackson Laboratory (C57BL/6-Tg [Pf4-cre] Q3Rsko/J). Cd36^fl/fl^ mice were purchased from The Jackson Laboratory (Cd36tm1.1Ijg/J) and crossed to Pf4-Cre mice and Hbb-Cre mice (C57BL/6-Tg [Hbb-Cre]12Kpe/J), respectively. Experiments were performed with male mice 4–5 weeks of age for mesenteric artery occlusion model and 10–12 weeks of age for carotid artery ligation and PPE model.

### Statistical analysis

Data are provided as arithmetic means ± standard error of mean (SEM), statistical analysis was performed using GraphPad Prism Version 7.0.5. Clinical data were initially evaluated to check the normality of the distribution using the Shapiro-Wilk test and subsequently the unpaired t-test with Welch’s correction or Mann-Whitney test was used. Comparison of two groups were performed using unpaired student’s t-test or multiple t-test. Comparisons of more than two groups were performed using a one- or two-way ANOVA with Sidak’s multiple-comparison. *P* values < 0.05 were considered to be statistically significant.

## Data availability

The original contributions presented in the study are included in the article, further inquiries can be directed to the corresponding author.

## ACKNOWLEDGEMENTS

We thank Dr. Maria Febbraio for providing *Cd36*^−/–^ mice and Julia Odendahl for performance of the PPE mouse model. We thank Martina Spelleken for excellent technical assistance. Special thanks goes to the CAi for the use of fluorescence microscopes. The study was supported by the *Deutsche Forschungsgemeinschaft* (DFG), grant number EL651/6-1 to M.E.; WA 3533/3-1 to MUW and Collaborative Research Centre TRR259 (Aortic Disease – Grant No. 397484323; projects A05 and A07 to N.G., to M.E. and H.S. Likewise, we thank the team of Prof. Dr. Hadi Al-Hasani (BK, AH, CH. and JS) and the IRTG 1902 at the HHU. We acknowledge the support of the Susanne-Bunnenberg-Stiftung at the Düsseldorf Heart Centre.

## AUTHOR CONTRIBUTIONS

ME and KJK designed the experiments, analyzed data and wrote the manuscript. HAH, AC and KJ provided *Cd36*^−/–^ and *Tsp-1*^−/–^ mice, respectively. MUW and HS provided AAA patient specimens. ANB carried out *Cd36*^−/–^ mice experiments under static conditions and analyzed TSP-1 plasma levels of AAA patients. ES-D, FR, KJK, TF performed whole blood studies of AAA patients. AE performed the FasL-ELISA of AAA patients and KJK performed all other experimental work. BEK, CR, NG, MK and CQ and ME handed out advices to perform experimental work and interpreted data. KK, IK and SP performed the carotid artery ligation of C57BL/6J mice and IK performed mesenteric artery occlusion model with cell specific (Cre) *Cd36*^−/–^ mice.

