## Supplemental data for "Crosstalk between thrombospondin-1 and CD36 modulates platelet–RBC interaction limiting thrombosis and abdominal aneurysm formation"

[1]Clinic of Vascular and Endovascular Surgery, Medical Faculty and University Hospital Düsseldorf, Heinrich-Heine-University, Moorenstraße 5, 40225 Düsseldorf, Germany. [2]Institute for Clinical Biochemistry and Pathobiochemistry, German Diabetes Center, Leibniz-Center for Diabetes Research at the Heinrich-Heine-University Düsseldorf, Medical Faculty, 40225 Düsseldorf, Germany. [3]Division of Cardiology, Pulmonology and Vascular Medicine, University Hospital and Medical Faculty, Heinrich-Heine-University, Düsseldorf, Germany. [4]Center of Thrombosis and Haemostasis (CTH), University Medical Center of the Johannes Gutenberg-University, Mainz. [5]German Center for Cardiovascular Research (DZHK), Partner Site Rhine Main, 55131 Mainz, Germany. [6]Department of Anesthesiology, Intensive Care and Pain Medicine, Experimental and Clinical Haemostasis, University of Muenster, 48149 Muenster, Germany; [7]OxProtect GmbH, 48149 Muenster, Germany. [8]INSERM U976, HIPI, Institut de Recherche Saint Louis, Paris, France; Université de Paris, France. [9]Cardiovascular Research Institute Düsseldorf (CARID), Medical Faculty, Heinrich-Heine-University, Düsseldorf, Germany.

Authorship note: MUW and ME are shared last authorship

### SUPPLEMENTAL METHODS

##### *Chemicals and Antibodies*

Platelets were activated by ADP (Sigma-Aldrich), CRP (collagen-related peptide, CambCol Laboratories, United Kingdom) and PAR4 activating peptide (AYPGKF, JPT Peptide Technologies). Human fibrinogen (Sigma-Aldrich), heparin (Ratiopharm), prostacyclin (Calbiochem), apyrase (Sigma-Aldrich, grade II, from potato), complete protease inhibitor cocktail (Roche) were all purchased. Accutase (catalog no. L11-007) PAA Laboratories GmbH. PE anti-human CD178 (FasL) (catalog no. 306406), BioLegend. Cy™5 annexin V (catalog no. 559933), PE mouse anti-human CD42a (catalog no. 558819) as well as PE mouse anti-human CD61 (catalog no. 555754), BD Biosciences. CD235a-FITC (catalog no. IM2212U) was from Beckman Coulter. PE Fas receptor antibody (CD95, catalog no. 305608) was from BioLegend and Fas protein (catalog no. 10217-H08H) from Sino Biological. PE anti-CD36 (catalog no. 529010) was from Thermo Fisher Technology. HRP-conjugated anti-mouse IgG (catalog no. NA931) was from GE Healthcare. Recombinant human DcR3 (TNFRSF6B Fc Chimera) was from R&D Systems (catalog no. 095407). Anti-CD36 mAb (catalog no. 10009893) from Cayman chemicals. Anti-TSP-1 mAb (catalog no. NB100-2059), Novus Biologicals. ReoPro® (abciximab) and Aggrastat® (tirofiban), Janssen Biologics B.V. Fas receptor antibody CD95 (catalog no. GTX13549), GeneTex. Recombinant annexin V inhibiting protein (catalog no. 556416), BD Biosciences. Native thromospondin-1 (TSP-1) protein (catalog no. 60522525UG), Merck and Elastase (catalog no. 39445-21-1), Sigma-Aldrich. Unconjugated CD235 anti-human (catalog no. NBP2-45024), Novus Biologicals. Unconjugated GPIX (catalog no. orb1672889), Biorbyt such as Alexa Fluor™ 488 goat anti-mouse IgG (catalog no. A11006), and Alexa Fluor™ 555 goat anti-rabbit IgG (catalog no. A21428), Invitrogen. IgG1 (catalog no. 554112) and IgM (catalog no. 555583) from BD Biosciences. All other reagents were of analytical grade.

***Human whole blood***

Fresh citrate-anticoagulated blood (105 mM $Na_3$-citrate, BD-Vacutainer®; Becton, Dickinson and company) was obtained from healthy volunteers aged between 18 and 70 years and AAA patients. AAA patients were compared to age-matched controls (AMCs, older than 60 years). (study number: 2018-140-KFogU). Human AAA blood samples were collected pre-surgery. Tissue samples were collected and obtained from our local Biobank at the Department of Vascular- and Endovascular Surgery at University Hospital Düsseldorf (study number: 5731R and 2018-248-FmB).

***Human platelet and plasma preparation***

Human blood fresh citrate-anticoagulated blood (105 mM $Na_3$-citrate, BD-Vacutainer®; Becton, Dickinson and company) was collected and centrifuged at 231 *g* for 10 min. The upper phase consisting of the platelet-rich plasma (PRP) was carefully transferred into phosphate buffered saline (PBS) pH 6.5 containing apyrase (2.5 U/mL) and 1 µM $PGE_1$. The tubes were centrifuged at 1000 *g* for 6 min and the obtained pellet was resuspended in tyrode's buffer (137 mM NaCl, 2.8 mM KCl, 12 mM $NaHCO_3$, 0.4 mM $NaH_2PO_4$, 5.5 mM glucose, pH 6.5). The final cell count was measured by a hematology analyzer (Sysmex KX-21N, Norderstedt, Germany) and adjusted according to the requirements of the experiment. For Additional plasma collection, the tubes were centrifuged, after the second centrifugation step and the extraction of the PRP, at 1500 *g* for 10 min at 4 °C. The platelet-free plasma (PFP) was collected and either stored on ice for immediate usage or stored at –70 °C for later experiments.

***Isolation of human red blood cells (RBCs)***

After PRP separation, the remaining blood was centrifuged at 800 *g* for 15 min in a closed syringe. The plasma was removed and used to prepare cell-free plasma. To separate RBCs from leukocytes the syringe was opened and the red blood cells were collected in a new tube. Red blood cells were washed three times with the fivefold volume of saline solution (154 mM) by centrifugation at 300 *g* for 10 min. RBCs were used immediately. The final cell count was measured by a hematology analyzer (Sysmex KX-21N, Norderstedt, Germany) and adjusted according to the requirements of the experiment.

***Platelet preparation and total blood cell count in mice***

Platelets were prepared as previously described (1, 2). Briefly, murine blood was collected from retro-orbital plexus in 300 µL heparin (20 U/mL in PBS) and total blood cell counts were analyzed by a hematology analyzer (Sysmex KX-21N, Norderstedt, Germany). The blood samples were centrifuged at 250 *g* for 5 min. To obtain PRP, the supernatant was centrifuged at 50 *g* for 6 min. Plasma samples were taken after an additional centrifugation step with 650 *g* for 5 min at RT. After discarding the buffy coat, RBCs were taken. Sedimented platelets were washed twice with tyrode's buffer (136 mM NaCl, 0.4 mM $Na_2HPO_4$, 2.7 mM KCl, 12 mM $NaHCO_3$, 0.1% glucose, 0.35% bovine serum albumin [BSA; pH 7.4]) supplemented with prostacyclin (0.5 µM) and apyrase (0.02 U/mL) at 650 *g* for 5 min. Before use, platelets were resuspended in the same tyrode's buffer supplemented with 1 mM $CaCl_2$.

***Thrombus formation under flow ex vivo using the flow chamber system***

Glass coverslips (24 × 60 mm) were coated with collagen (200 µg/mL; Horm®, Takeda Pharmaceutical), TSP-1 protein (100 µg/mL; catalog no. 605225-25UG) or collagen/TSP-1 incubated overnight at 4 °C and blocked with 1% BSA for at least 1 hour at RT. Whole blood or recomposed blood was perfused through the chamber (50 µm × 5 mm). Human or murine platelets as well as PRP, PFP plasma and RBCs were isolated according to the standard protocols above. The cell counts were adjusted to $2 \times 10^5$ platelets/µL and $4 \times 10^6$ RBCs/µL for human samples or $3 \times 10^5$ platelets/µL and $4 \times 10^6$ RBCs /µL for mouse samples using PFP plasma. Murine blood components were reassembled as indicated in the individual experiment. Blocking studies in human whole blood were performed using an anti-TSP-1 mAb (2 µg/mL; catalog no. NB100-2059), an anti-CD36 mAb (2 µg/mL; catalog no. 10009893) for 10 min, prior to flow chamber experiments. For single cell type blocking experiments, either RBCs or platelets were incubated with an anti-CD36 mAb (2 µg/mL; catalog no. 10009893), an anti-TSP-1 mAb (2 µg/mL; catalog no. NB100-2059) or respective IgG control antibody (2 µg/mL; catalog no. 554121) for 10 min. Subsequently, RBCs were washed and used for the experiments. In every experiment, Mepacrin (10 µM; human experiments) or DyLight 488-conjugated anti-mouse (GP) Ibβ antibody (2 µg/mL; murine experiments) was added and incubated for 10 min. After incubation, the blood was perfused over a collagen-coated surface at a shear rate of 1,000 $s^{-1}$ or 1,700 $s^{-1}$ using a pulse-free electric pump. After a predefined timespan (3 min), the blood perfusion was stopped and the flow chamber was perfused with tyrode's buffer at a shear rate of 1,000 $s^{-1}$. Five pictures from different areas were taken (400-fold total magnification; Axio Observer.D1, Carl Zeiss). The surface coverage was analyzed using the ImageJ-win64 software. Where indicated cells were resolved by Accutase® treatment for 15 min (120 µL per coverslip; solution with protease and collagenolytic activity) to detach and solve adherent and aggregated cells and analyzed by flow cytometry.

The cell suspension for PS exposure analysis was incubated in the dark for 15 min with an antibody mix containing CD42a-PE (catalog no. 558819), CD235a-FITC (catalog no. B49206) and annexin V-Cy5 (catalog no. 559933). In mice, isolated platelets were incubated with annexin V-Cy5 alone. The cell suspension for TSP-1 binding was incubated with an anti-TSP-1 mAb (catalog no. NB100-2059) for 30 min and subsequently incubated with Alexa Fluor™ 488 goat anti-mouse IgG (catalog no. A11001) for further 30 min in the dark, respectively. The cell suspension for CD36 binding was incubated with an antibody mix containing CD235-FITC and CD36-PE (catalog no. A15793) for 15 min in the dark. The incubation was stopped by adding 300 µL binding buffer or PBS and the samples were analyzed using a BD FACSCalibur™ (BD Bioscience).

##### *Adhesion study*

Adhesion experiments on immobilized FasR protein and fibrinogen were performed with isolated human platelets and Chinese hamster ovary cells (CHO cells) as well as integrin $\alpha_{IIb}\beta_3$ transfected into Chinese hamster ovary cells (A5 CHO cells). Glass coverslips (24 × 60 mm) were either coated with FasR protein (50 µg/mL) or fibrinogen (100 µg/mL) at a defined area

(10 × 10 mm); incubated in humidity chambers at 4 °C overnight. Coverslips were blocked with 1% BSA for at least 1 h at RT. Resting or ADP (10 μM)-stimulated platelets (4 × $10^3$/μL) were pretreated with the inhibitors hDcR3 (10 μg/mL, recombinant human decoy receptor 3 protein), tirofiban (1 μg/mL) and abciximab (10 μg/2 million platelets) for 10 min and then allowed to adhere to the coated coverslips for 60 min at RT. IgG-FC (10 μg/mL) peptide served as control. CHO and A5 CHO cells were allowed to adhere to the coated coverslips for 30 min. Afterwards, cover slips were rinsed two times with PBS to wash off unbound platelets and CHO cells. Adherent cells were fixated immediately with 4% paraformaldehyde (PFA) and covered with the mounting medium (Aquatex®). Five DIC images from different areas were taken (platelets: 400-fold total magnification; CHO cells: 100-fold total magnification; Axio Observer.D1, Carl Zeiss). The total number of adherent platelets was counted using ImageJ-win64 software.

***Platelet aggregation***

Impedance measurements were conducted as describes elsewhere (3). Briefly, impedance measurement of platelets supplemented with plasma in the absence or presence of RBCs as well as whole blood of human or mice was performed compared to NaCl (0.9% saline) buffer using Chrono-Log dual channel lumi-aggregometer® (model 700) at 37 °C stirring at 1,000 rpm. Human blood was collected into Vacutainer® sodium citrate tubes and murine blood in ACD buffer. Suspensions of murine platelets in the absence or presence of RBCs as well as whole blood of knockout mice were stimulated with collagen (10 μg/mL collagen in whole blood or 20 μg/mL collagen in samples with platelets in the absence or presence of RBCs) in the presence of fibrinogen (70 μg/mL). Respective human samples were stimulated with 10 μg/mL collagen without fibrinogen. Isolated human and murine platelets and RBCs were reassembled with PFP plasma to a concentration of 4 × $10^6$ RBCs/μL and 2 × $10^5$ platelets/μL (human) and 4 × $10^6$ RBCs/μL and 3 × $10^5$ platelets/μL (mouse). Where indicated, anti-TSP-1 mAb was used in a concentration of 2 μg/mL for 10 min.

***Flow cytometry***

Isolated platelets and RBCs were adjusted in equal concentrations for flow cytometric analysis. Whole blood (heparinized mouse and fresh citrate-anticoagulated human blood) was diluted 1:10 in tyrode's buffer. For flow cytometry experiments, all antibodies were diluted 1:10 in a total reaction volume of 30 µL, incubated for 15 min at RT in the dark, and stimulated with indicated agonist. Staining was stopped by addition of 300 µL PBS and subsequently analyzed on a FACSCalibur™ (BD Bioscience). For annexin V-Cy5 staining, binding buffer (10 mM HEPES, 140 mM NaCl, 2.5 mM $CaCl_2$, pH 7.4) was used (4). CD42-PE served as platelet specific marker, while CD235-FITC was used as human RBC specific cell marker. For TSP-1 binding, an anti-TSP-1 mAb was preincubated for 30 min at RT and subsequently incubated with Alexa Fluor™ 488 goat anti-mouse IgG (catalog no. A11001) for further 30 min in the dark. Where indicated, cells were pre-treated with ReoPro® (abciximab, 10 µg/2 million cells), Aggrastat® (tirofiban, 1 µg/mL) anti-FasR mAb (CD95, 10 µg/mL), hDcR3 (10 µg/mL), anti-TSP-1 mAb (2 µg/mL) or anti-CD36 mAb (2 µg/mL) for 15 min at RT. For analysis, platelets were gated using their specific forward scatter (FSC) and side scatter (SSC) profile and/or with their specific cell marker profile. Externalization of Fas ligand (FasL-PE), PS exposure, CD36 (CD36-PE), FasR (FasR-PE) and inactive integrin $\alpha_{IIb}\beta_3$ (CD61-PE; $\beta_3$ integrin subunit) on activated and non-activated platelets was determined by flow cytometry in the absence or presence of RBCs. The inactive form of integrin $\alpha_{IIb}\beta_3$ in murine samples was stained with CD61-FITC (catalog no. M031-1, Emfret Analytics). For the analysis of CD45- and Ly6G-platelet aggregates, specific antibodies were used (Emfret Analytics, GPIb/CD42b, Xia.G5-PE; BD Biosciences, anti-mouse CD45-APC, catalog no. 559864 and BioLegend, anti-mouse Ly-6G-APC, catalog no. 127614).

***Western blot***

Briefly, Platelets and RBCs were isolated following the standard protocols. Cell counts were adjusted according to the requirements of the experiment. Where indicated, samples were incubated with CRP for 5 min and for further 10 min at 37 °C in the absence and presence of

RBCs. TSP-1 was determined in platelet lysates and releasates. Each sample had a total number of 40 × $10^6$ platelets and/or 200 × $10^6$ RBCs, which were sedimented by centrifugation and lysates were prepared, separated on SDS-polyacrylamide gel and transferred onto nitrocellulose blotting membranes. Subsequently, the membrane was blocked with 5% BSA or 5% (w/v) powdered skim milk in TBS-T (tris-buffered saline with 0.1% Tween 20) for 60 min and probed with an anti-TSP-1 mAb (catalog no. NB100-2059, 1:1000). The antibody incubations were performed at 4 °C overnight. On the next day, the membrane was washed three times with TBS-T, and incubated with the corresponding horseradish peroxidase (HRP)-conjugated secondary anti-rabbit antibodies in 5% powdered skim milk in TBS-T (1:2500, catalog no. NA934) for 1 h at RT. For visualizing protein bands the Vilber Fusion-FX6-EDGE V.070 imaging system was used.

***Quantification of FasL and TSP-1 by ELISA***

Circulating FasL, CD36 and TSP-1 in plasma samples of AAA patients and age-matched controls (AMCs) were determined following the manufacturer's protocol for human specific FasL ELISA (catalog no. ab100515, abcam®), CD36 ELISA (catalog no. EK0700, Booster Biological Technology) and human specific TSP-1 ELISA (catalog no. ab193716, abcam®).

***Histology***

For the analysis of platelets, RBCs and TSP-1 in samples of aortic vessel walls, 5-micron sections of paraffin-embedded tissue from intraluminal thrombi of AAA patients and occluding thrombi of patients undergoing thrombectomy (peripheral thrombi) were prepared. For immunofluorescence staining of platelets, RBCs and TSP-1 paraffin-embedded sections were stained for platelets with an anti-GPIX mAb (rabbit anti-mouse/human GPIX, Biorbyt, catalog no. orb167288, 1:100, 20 µg/mL) followed by an Alexa Fluor™ 555 labeled secondary antibody (goat anti-rabbit, Invitrogen, 1:100). For RBC staining anti-CD235 mAb (rat anti-human CD235, Novus Biologicals, catalog no. NBP2-45024 1:200, 0.5 µg/mL) followed by an Alexa Fluor™ 488 labeled secondary antibody (goat anti-mouse, Invitrogen, 1:100). For TSP-1 staining anti-TSP-1 mAb (rabbit anti-human TSP-1, Novus Biologicals, 1:100, 20 µg/mL) followed by an Alexa Fluor ™ 555 labeled secondary antibody (goat anti-rabbit, Invitrogen, 1:100) were used. Nuclei were identified using DNA staining with 4, 6 diamidino-2-phenylindole dihydrochloride (DAPI, Roche, 1:3000). IgG served as control and were used in the same concentration as the depending primary antibodies, respectively. Additionally, fluorescence emission at 488 nm showed autofluorescence of red blood cells in IgG controls and autofluorescence of elastic lamina in vessel wall samples. Aortic tissue samples of mice were stained with purified rat anti-mouse GPIbα (CD42b, 10 µg/mL, Emfret Analytics, catalog no. M042-0) and incubated at 4 °C overnight. On the next day, the appropriate secondary antibody, eBioscience™ streptavidin eFluor™ 660 conjugate (10 µg/mL, Invitrogen™, catalog no. 50-4317-80) were applied to the tissues at RT for 1h. In case of platelet staining with CD42b, an intermediate step of biotinylation was used to enhance the fluorescence signal. Samples were analysed using confocal microscopy (LSM 710, Carl Zeiss, Germany).

**Intravital microscopy of thrombus formation in mesenteric arterioles injured with $FeCl_3$**

Intravital microscopy was performed as described previously (6). Male mice (4–5 week of age) were anesthetized with Ketamin (Ketavet®, Pfizer, 100 mg/kg) and Xylazin (2% Bernburg, medistar, 5 mg/kg) by intraperitoneal (i.p.) injection. Thrombus formation was made visible by intravenous (i.v.) injection of DCF (4.48 mM, 100 µL, Sigma-Aldrich) and observed with a fluorescence microscope (10-fold, PH1, Zeiss, Axio Observer, Germany). After a midline abdominal incision, the mesentery was exteriorized and a filter paper saturated with 20% $FeCl_3$ was used to injure arterioles by topical application for 20 seconds. Time until full occlusion of the vessel (when blood flow had stopped for 60 seconds) was measured. Experiments were stopped after 40 min. Euthanasia was performed by cervical dislocation.

***Carotid artery ligation model***

Carotid artery ligation was performed in male mice 12 weeks of age at the Center for Thrombosis and Hemostasis Mainz (CTH; *Tsp-1*$^{-/-}$ platelets) and at the University hospital

Düsseldorf (UKD; *Cd36*$^{-/-}$ platelets) as described elsewhere (5). Platelets from *Tsp-1*$^{-/-}$ and *Cd36*$^{-/-}$ donor mice were stained with Rhodamine B or CellTracker™ Red CMTPX (Invitrogen™) according to the manufacturer's guidelines. Either C57BL/6J mice or WT littermates (*Cd36*$^{+/+}$) mice were anaesthetized as described above and put on a heating pad. The right common carotid artery was prepared, and after intravenous injection of fluorescently labeled platelets, a film was taken using a DM6FS microscope (Leica Microsystems, Wetzlar, Germany) or BX51WI microscope (Olympus, Hamburg, Germany). Afterwards, the carotid artery was ligated vigorously for 5 min, thus inducing a vascular injury. The interaction of fluorescent platelets with the injured vessel wall was visualized 20 min after ligation by *in vivo* video microscopy. Euthanasia was performed by cervical dislocation.

***Porcine pancreatic elastase infusion model***

10 to 12-week-old C57BL/6J (*Tsp-1*$^{+/+}$) or *Tsp-1*$^{-/-}$ mice were used. To induce experimental AAA, mice were anesthetized with 2–3% isoflurane and received a locally subcutaneous (s.c.) injection of buprenorphine (0.1 mg/kg) (Temgesic, Eumedica) 30 min before surgery. After median laparotomy, the proximal and distal infrarenal aorta were temporarily ligated to create an aortotomy above the iliac bifurcation. A catheter was inserted into the distal end of the aortotomy and used to infuse the aorta with sterile isotonic saline (NaCl, 0.9%) containing type I Porcine Pancreatic Elastase (2.5 U/mL) (Sigma-Aldrich, catalog no. E1250) at 120 mm Hg for 5 min. The vessels in sham mice were infused with sodium chloride only. After removal of the infusion catheter, the aortotomy was closed without constriction of the aortal lumen. Finally, the abdomen was closed. For pain relief, all animals received Buprenorphine s.c. every 6 h during the daylight phase for 3 days and in the dark phase via drinking water (0.3 µg/mL). At the endpoint at day 28 mice were euthanized under deep isoflurane anesthesia followed by cervical dislocation.

***Ultrasound imaging***

Prior PPE surgery (baseline) and at days 3, 7, 14, 21 and 28 post-surgery, maximal aortic diameters at the aneurysm site were measured using ultrasound. Mice were anaesthetized with 2–3% isoflurane, placed on a 37 °C heated plate and ultrasound imaging was performed using a Vevo 2100® High-Resolution In Vivo Micro-Imaging System (VisualSonics). Inner diameter-measurements were obtained following a standardized imaging algorithm with longitudinal B-Mode images during the systolic phase.

SUPPLEMENTAL FIGURES

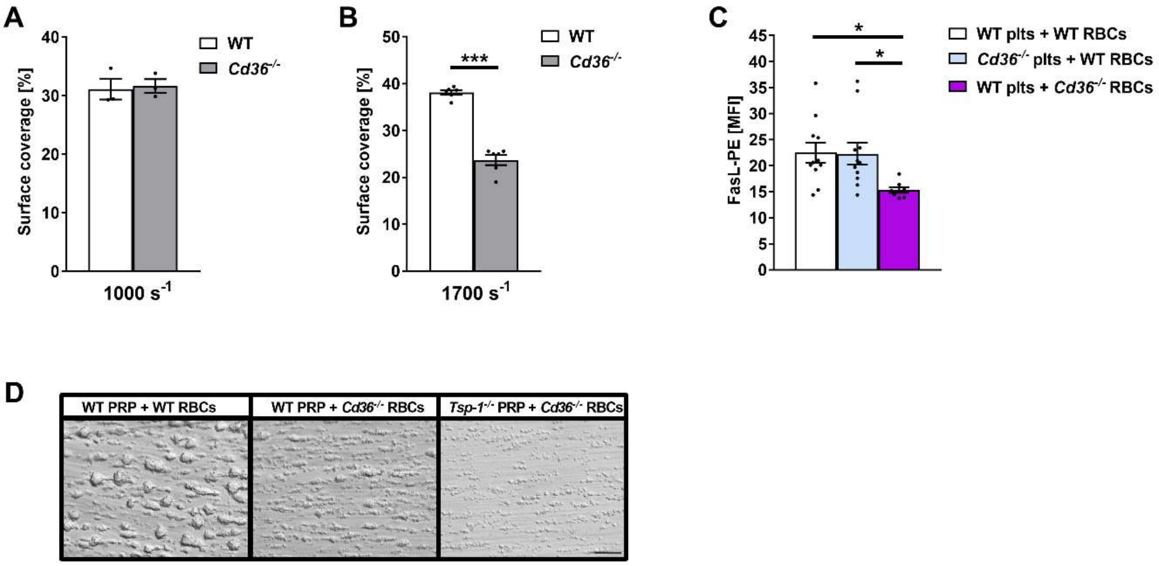


**Figure S1. Impaired collagen-dependent thrombus formation of *Cd36*$^{-/-}$ mice under high shear-rates and reduced FasL externalization on platelets after incubation with *Cd36*$^{-/-}$ RBCs.** (**A** and **B**) Surface coverage of thrombus formation on collagen (200 µg/mL) at a shear rate of (**A**) 1,000 s$^{-1}$ and (**B**) 1,700 s$^{-1}$ using whole blood from WT and *Cd36*$^{-/-}$ mice ($n$ = 3–6). (**C**) Externalization of FasL on the surface of WT and *Cd36*$^{-/-}$ platelets in the presence of either WT or *Cd36*$^{-/-}$ RBCs was determined by flow cytometry ($n$ = 12). (**D**) Representative images of thrombus formation of either WT or *Tsp-1*$^{-/-}$ platelets incubated with WT or *Cd36*$^{-/-}$ RBCs ($n$ = 3). Data are represented as mean values ± SEM. *$P$ < 0.05; ***$P$ < 0.001 tested by student's t-test (**A** and **B**) and one-way ANOVA Sidak's multiple comparison test. WT, wild-type; plts, platelets; RBCs, red blood cells; MFI, mean fluorescence intensity.

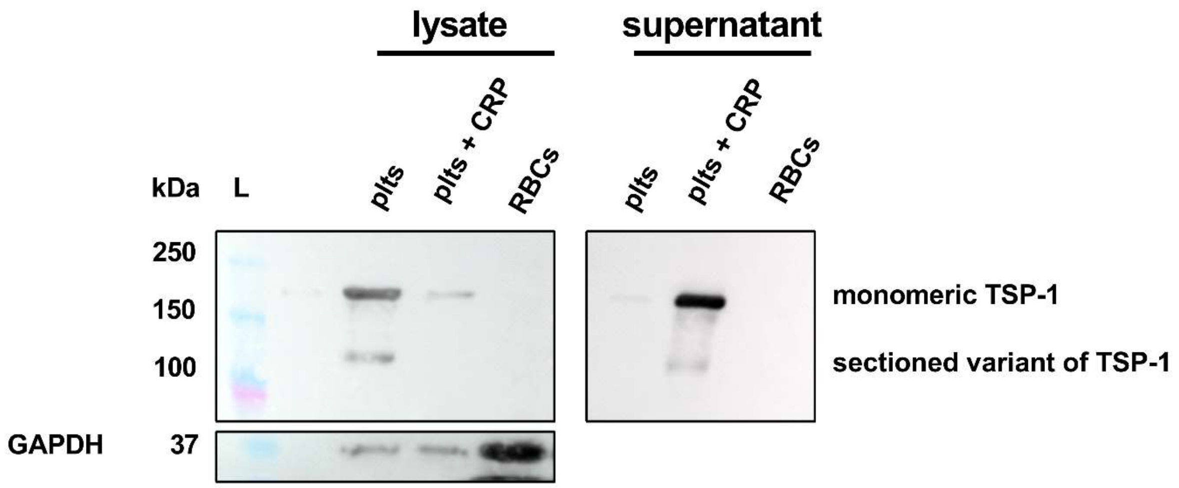


**Figure S2. TSP-1 protein abundance of platelets and RBCs in healthy human volunteers.** Human platelets were activated with CRP (5 µg/mL) for 5 min at 37 °C or left under resting conditions in the absence or presence of RBCs. Subsequently, lysates and supernatants were generated and western blot analysis was performed (*n* = 4). GAPDH served as loading control. Plts, platelets; RBCs, red blood cells; CRP, collagen-related peptide; L, ladder; TSP-1, thrombospondin-1.

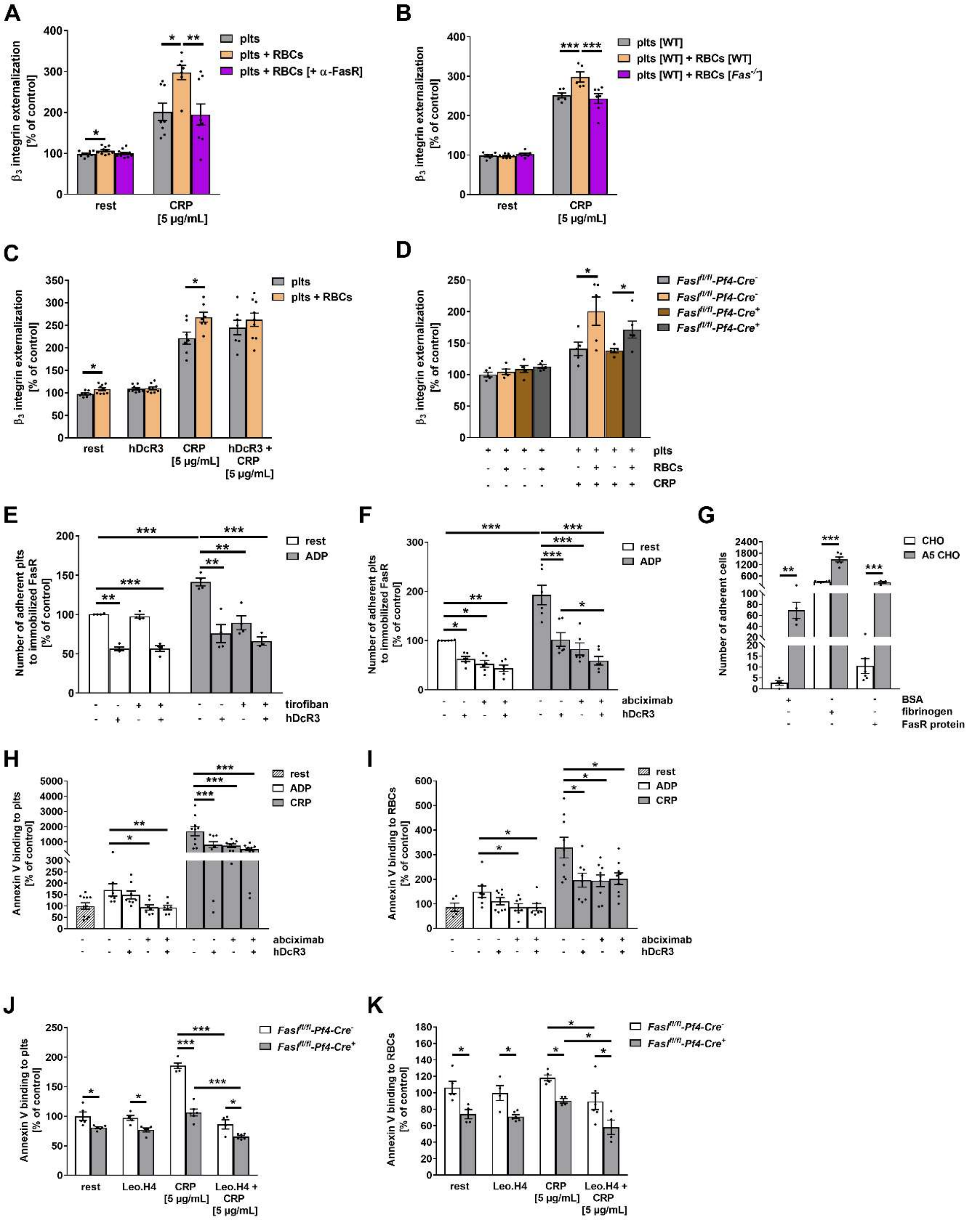


**Figure S3. Integrin $\alpha_{IIb}\beta_3$ serves as another ligand on the platelet membrane important for FasR activation on RBCs.** (**A**–**D**) Externalization of integrin $\beta_3$ subunit on the surface of human and murine (*Fas*$^{-/-}$ and *Fasl*$^{fl/fl}$-*Pf4-Cre*) platelets in the absence or presence of RBCs after stimulation with CRP (5 µg/mL) was determined by flow cytometry. (**A** and **C**) Human RBCs were treated with an anti-FasR mAb (**A**, 10 µg/mL, *n* = 7-9) and human platelets with hDcR3 (**C**, 10 µg/mL, *n* = 7-9). IgG treatment served as control. (**B**) Integrin $\beta_3$ subunit externalization of WT platelets alone and with either WT or *Fas*$^{-/-}$ platelets (*n* = 6-7) (**D**) Either WT (*Fasl*$^{fl/fl}$-*Pf4-Cre*$^{-}$) or FasL-deficient (*Fasl*$^{fl/fl}$-*Pf4-Cre*$^{+}$) platelets were incubated with RBCs from WT mice (*n* = 5). (**E** and **F**) Quantification of adherent resting and ADP (10 µM)-stimulated platelets to immobilized recombinant FasR protein (50 µg/mL) in the absence and presence of

FasL inhibitor (hDcR3, 10 µg/mL) and integrin $\alpha_{IIb}\beta_3$ inhibitors (**E**, tirofiban, 1 µg/mL; **F**, abciximab, 10 µg/2 million cells) (*n* = 4–6). Control samples were incubated with IgG-Fc (*n* = 4–6). (**G**) Adhesion of CHO and A5 CHO cells to immobilized recombinant FasR protein and fibrinogen (100 µg/mL) (*n* = 5). BSA (1%) served as control (*n* = 5). (**H–K**) Annexin V binding to either human (**H** and **I**, *n* = 9) or murine (**J** and **K**, *n* = 6) platelets and RBCs, after stimulation with indicated agonists in the absence and presence of FasL inhibitor (hDcR3, 10 µg/mL) and integrin $\alpha_{IIb}\beta_3$ inhibitor (abciximab, 10 µg/2 million cells; mouse, Leo.H4, 5 µg/mL) was determined by flow cytometry. Depicted as percent-gated cells normalized to controls. Data are represented as mean values ± SEM. **P* < 0.05; ** *P* < 0.01;*** *P* < 0.001 tested by two-way ANOVA (**A, B, D, J, K**), multiple t-test (**C, G**) and one-way ANOVA (**E, F, H, I**) with Sidak's multiple comparison test. Plts, platelets; RBC, red blood cells; WT, wild-type; rest, resting; CRP, collagen-related peptide; ADP, adenosine diphosphate; abciximab, tirofiban, Leo.H4, $\alpha_{IIb}\beta_3$ inhibitor; hDcR3 (human decoy receptor 3), FasL inhibitor; BSA, bovine serum albumin; CHO cells, Chinese hamster ovary cells; A5 CHO cells, integrin $\alpha_{IIb}\beta_3$ transfected into Chinese hamster ovary cells.

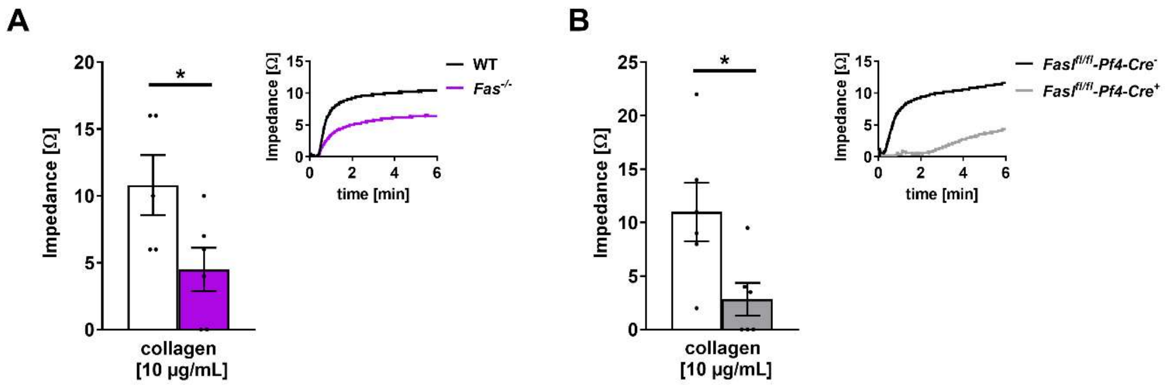


**Figure S4. Genetic deletion of FasR and FasL results in impaired collagen-dependent platelet aggregation.** (**A** and **B**) Impedance measurements of murine whole blood after stimulation with collagen was performed. Representative aggregation curves are shown. (**A**) Whole blood of *Fas*$^{-/-}$ mice was stimulation with collagen (10 µg/mL) ($n$ = 6). WT mice served as control ($n$ = 6). (**B**) Whole blood of *Fasl*$^{fl/fl}$*-Pf4-Cre*$^{+}$ mice was stimulation with collagen (10 µg/mL) ($n$ = 6). WT mice (*Fasl*$^{fl/fl}$*-Pf4-Cre*$^{-}$) served as controls ($n$ = 6). Data are represented as mean values $\pm$ SEM. *$P$ < 0.05 tested by student's t-test (**A, B**). WT, wild-type; CRP, collagen-related peptide.

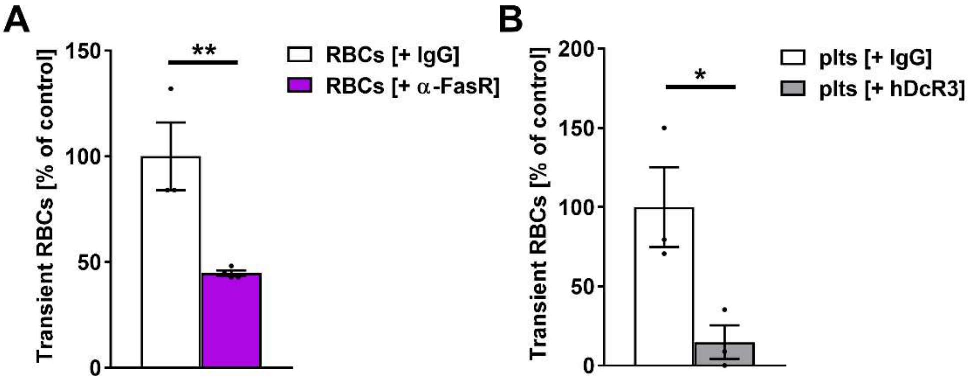


**Figure S5. Antibody-mediated inhibition of FasL and FasR reduces RBC recruitment to collagen-adherent platelets.** (**A** and **B**) Experiments with cell type-specific antibody treatment of either RBCs with an anti-FasR mAb (**A**, 10 µg/mL) or collagen-adherent platelets with hDcR3 (**B**, 10 µg/mL) ($n$ = 3–4). IgG treatment served as control ($n$ = 3–4). Data are represented as mean values ± SEM. *$P$ < 0.05; **$P$ < 0.01 tested by student's t-test (**A, B**). Plts, platelets; RBCs, red blood cells.

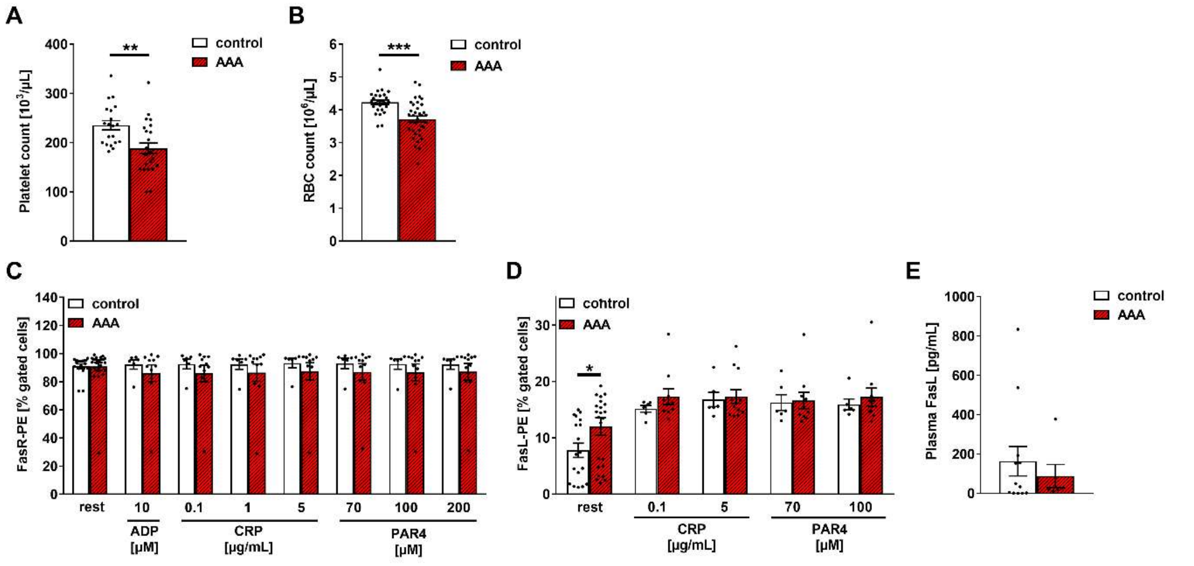


**Figure S6. Reduced blood cell count and increased FasL on platelets in AAA patients.** (**A**) Platelet count in AAA patients compared to age-matched controls (controls, *n* = 20; AAAs, *n* = 24). (**B**) RBC count in AAA patients compared to age-matched controls (controls, *n* = 26; AAAs, *n* = 35). (**C**) FasR externalization on RBCs after stimulation with indicated agonists (controls, *n* = 5–18; AAAs, *n* = 8–25). (**D**) Externalization of FasL on the platelet surface of AAA patients compared to age-matched controls (controls, *n* = 6–18; AAAs, *n* = 10–24). (**E**) Analysis of soluble FasL in plasma of AAA patients compared to age-matched controls (controls, *n* = 12; AAAs, *n* = 6). Data are represented as mean values ± SEM. **P* < 0.05; ***P* < 0.01; ****P* < 0.001 tested by unpaired t-test with Welch's correction (**A, B**), multiple t-test (**C, D**) and Mann-Whitney test (**E**). Plts, platelets; RBC, red blood cells; rest, resting; AAA, abdominal aortic aneurysm; control, age-matched controls; ADP, adenosine diphosphate; CRP, collagen related peptide; PAR4, protease-activated receptor 4 activation peptide.

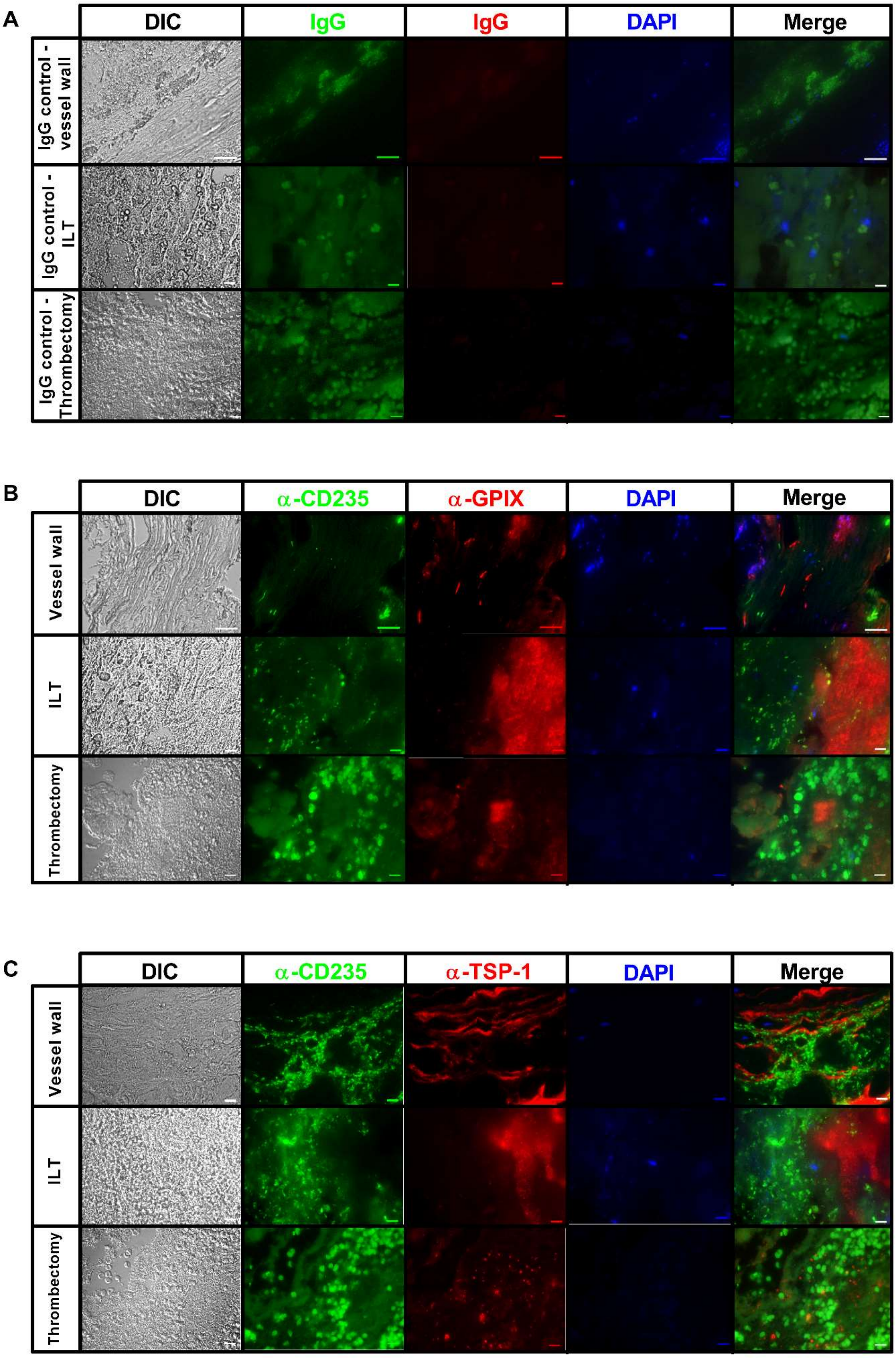
A
DIC
IgG
IgG
DAPI
Merge
IgG control - vessel wall
IgG control - ILT
IgG control - Thrombectomy
B
DIC
α-CD235
α-GPIX
DAPI
Merge
Vessel wall
ILT
Thrombectomy
C
DIC
α-CD235
α-TSP-1
DAPI
Merge
Vessel wall
ILT
Thrombectomy

**Figure S7. Immunofluorescence staining of AAA aortic vessel walls, ILT and thrombi from patients who underwent thrombectomy.** (**A**–**C**) Paraffin-embedded sections (5 µm) of aortic walls, the ILT from AAA patients and thrombi of patients who underwent thrombectomy (peripheral thrombi, lower panel) were stained with an anti-GPIX mAb (20 µg/mL) to detect platelets, an anti-CD235 mAb (0.5 µg/mL) to detect RBCs, and an anti-TSP-1 mAb (20 µg/mL) to detect TSP-1. IgG treatment served as control (same concentration as respective primary antibody). Additionally, fluorescence emission at 488 nm showed autofluorescence of red blood cells in IgG-treated controls and autofluorescence of elastic lamina in vessel walls. 4',6-diamidino-2-phenylindole (DAPI) was used as nucleus staining. Representative DIC and fluorescence images with 400-fold (**A** and **B**, upper panels, scale bar: 50 µm) and 1000-fold (**A and B,** middle and lower panels and **C** all images, Scale bar: 10 µm) magnification are shown. AAA, abdominal aortic aneurysm; ILT, intraluminal thrombus.

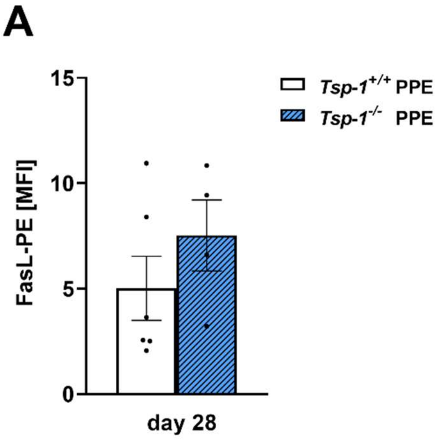


**Figure S8. FasL on platelets is unaltered in PPE-operated *Tsp-1*$^{-/-}$ mice 28 days post-surgery.** (**A**) Externalization of FasL on the platelet surface of PPE-operated *Tsp-1*$^{-/-}$ mice at day 28 post-surgery ($n$ = 4). *Tsp-1*$^{+/+}$ served as control ($n$ = 6). Tsp-1, thrombospondin-1.

#### Supplemental Videos

**Videos S1–4. *Ex vivo* tethering of human RBCs on collagen-adherent platelets under arterial shear rate (TSP-1–CD36 axis).** Isolated human platelets were stimulated with ADP (10 µM) and allowed to adhere on collagen (200 µg/mL) coated coverslips for 10 min. Representative videos of tethering RBCs on collagen-adherent platelets under flow using a shear rate of 1,700 $s^{-1}$. Collagen-adherent platelets or RBCs were treated with a blocking anti-CD36 mAb (2 µg/mL; platelets, Supplemental Video 1 and RBCs, Supplemental Video 3) or with corresponding IgG (2 µg/mL; platelets, Supplemental Video 2 and RBCs, Supplemental Video 4). Scale bar: 50 µm.

**Videos S5 and S6. *Ex vivo* tethering of human RBCs on collagen-adherent platelets under arterial shear rate (TSP-1–CD36 axis).** Isolated human platelets were stimulated with ADP (10 µM) and allowed to adhere on collagen (200 µg/mL) coated coverslips for 10 min. Representative videos of tethering RBCs on collagen-adherent platelets under flow using a shear rate of 1,700 $s^{-1}$. Collagen-adherent platelets were treated with a blocking anti-TSP-1 mAb (2 µg/mL; Supplemental Video 5) or with corresponding IgG (2 µg/mL; Supplemental Video 6). Scale bar: 50 µm.

**Videos S7–10. *Ex vivo* tethering of human RBCs on collagen-adherent platelets under arterial shear rate (FasL–FasR axis).** Isolated human platelets were stimulated with ADP (10 µM) and allowed to adhere on collagen (200 µg/mL) coated coverslips for 10 min. Representative videos of tethering RBCs on collagen-adherent platelets under flow using a shear rate of 1,700 $s^{-1}$. Collagen-adherent platelets were treated with hDcR3 (10 µg/mL; Supplemental Video 7) or with corresponding IgG-FC protein (10 µg/mL; Supplemental Video 8). RBCs were treated with a blocking anti-FasR mAb (10 µg/mL; Supplemental Video 9) or with corresponding IgG (10 µg/mL; Supplemental Video 10). Scale bar: 50 µm.

**Videos S11–14. *In vivo* stable platelet adhesion and thrombus formation after injury of carotid artery in C57BL/6J mice or WT littermates.** The right common carotid arteries of WT mice (C57BL/6J, *Tsp-1*$^{+/+}$) or WT littermates (*Cd36*$^{+/+}$) were prepared, intravenous injection of fluorescently labeled platelets and vessel injury (ligation for 5 min) was performed. Platelets from *Cd36*$^{-/-}$ (Supplemental Video 11) and *Tsp-1*$^{-/-}$ (Supplemental Video 13) donor mice and their corresponding controls (*Cd36*$^{+/+}$, Supplemental Video 12 and *Tsp-1*$^{+/+}$, Supplemental Video 14) were fluorescently labelled with rhodamine B or CellTracker™ Red CMTPX (Invitrogen™) according to the manufacturer's guidelines. The interaction of fluorescent platelets with the injured vessel wall was visualized 20 min after ligation by *in vivo* intravital microscopy. Videos showing carotid artery vessel wall, which is outlined using dotted lines.

**Videos S15–18. *In vivo* platelet adhesion and thrombus formation after injury of mesenteric arterioles in C*d36$^{fl/fl}$-Pf4-Cre and Cd36$^{fl/fl}$-Hbb-Cre mice*.** Mesenteric arterioles of *Cd36$^{fl/fl}$-Pf4-Cre$^{+}$* (Supplemental Video 15), *Cd36$^{fl/fl}$-Pf4-Cre$^{–}$* (Supplemental Video 16), and *Cd36$^{fl/fl}$-Hbb-Cre$^{+}$* (Supplemental Video 17) *and Cd36$^{fl/fl}$-Hbb-Cre$^{–}$* mice (Supplemental Video 18) were injured by topical application of 20% $FeCl_3$. Platelets were fluorescently labelled by intravenous injection of DCF and initial adhesion of platelets and thrombus formation was observed until full occlusion of the vessel using a fluorescent microscope (Zeiss Axio-Observer, objective: 10 x).
